# Pseudohyphal growth of the emerging pathogen *Candida auris* is triggered by genotoxic stress through the S phase checkpoint

**DOI:** 10.1101/832931

**Authors:** Gustavo Bravo Ruiz, Zoe K. Ross, Neil A.R. Gow, Alexander Lorenz

## Abstract

The morphogenetic switching between yeast cells and filaments (true hyphae and pseudohyphae) is a key cellular feature required for full virulence in many polymorphic fungal pathogens, such as *Candida albicans*. In the recently emerged yeast pathogen *Candida auris*, occasional elongation of cells has been reported. However, environmental conditions and genetic triggers for filament formation have remained elusive. Here, we report that induction of DNA damage and perturbation of replication forks by treatment with genotoxins, such as hydroxyurea, methyl methanesulfonate, and the clinically relevant fungistatic 5-fluorocytosine, causes filamentation in *C. auris*. The filaments formed were characteristic of pseudohyphae and not parallel-sided true hyphae. Pseudohyphal growth is apparently signalled through the S phase checkpoint and, interestingly, is Tup1-independent in *C. auris*. Intriguingly, the morphogenetic switching capability is strain-specific in *C. auris*, highlighting the heterogenous nature of the species as a whole.

**IMPORTANCE:** *Candida auris* is a newly emerged fungal pathogen of humans. This species was first reported in 2009, when it was identified in an ear infection of a patient in Japan. However, despite intense interest in this organism as an often multidrug-resistant fungus there is little knowledge about its cellular biology. During infection of human patients, fungi are able to change cell shape from ellipsoidal yeast cells to elongated filaments to adapt to various conditions within the host organism. There are different types of filaments, which are triggered by reactions to different cues. *Candida auris* fails to form filaments when exposed to triggers that stimulate yeast-filament morphogenesis in other fungi. Here, we show that it does form filaments when its DNA is damaged. These conditions might arise when *Candida auris* cells interact with host immune cells, or growing in certain host tissues (kidney, bladder), or during treatment with antifungal drugs.

## INTRODUCTION

The emergence of new multidrug resistance pathogens poses a recurrent global threat to healthcare settings. This is the case for the fungus *Candida auris* discovered as a new human pathogen only ten years ago (1), albeit a retrospective review of strain collections dated the first case back to 1996 (2). Since its first identification, *C. auris* has been found across all continents causing clonal outbreaks in hospital settings (3). It shows mortality rates in systemic disease close to 50 % (4), is one of the most drug-resistant yeast pathogens (5), and it has been described as a skin colonizer able to undergo nosocomial spread (6), thus becoming a major concern for medical mycology.

Due to its recent emergence, we are largely ignorant about its general biological traits. This lack of fundamental understanding about the origin (7, 8) and the life cycle of *C. auris* impedes our capacity to explain its sudden emergence, rapid global spread, and unique phenotypic characteristics. An example is the lack of information about its ability to undergo morphogenetic switches as described for other fungi. In fungi, a morphogenetic switch enables cells to change from growing as unicellular yeasts to pseudohyphae or true hyphae and can be triggered by a multitude of environmental factors, such as nutrient limitation, temperature, and pH changes (reviewed in (9–12)). Filamentous growth allows the exploration of new environments and is considered a virulence trait in pathogenic fungi (reviewed in (13, 14)). However, most cues causing filamentation in the best-studied and only distantly-related pathogen *C. albicans* do not induce filamentous growth in *C. auris* (15).

In contrast to true hyphae, pseudohyphal growth has been associated with delay in cell cycle progression and the subsequent extension of the apical growth period (16). Indeed, it has been demonstrated that drugs causing genotoxic stress, such as hydroxyurea (HU) or methyl methanesulfonate (MMS), trigger S phase arrest via a cell cycle checkpoint (reviewed in (17)); this results in pseudohyphal growth in *C. albicans* and *S. cerevisiae* (18, 19). The S phase checkpoint is a surveillance system, which responds to DNA damage or DNA replication fork arrest, and involves the sensor kinases Mec1 and Tel1, the mediator proteins Rad9 and Mrc1, and an effector kinase, Rad53. Rad9 acts as the main mediator for DNA damage response, whereas Mrc1 functions as DNA replication arrest responder (17). Once activated, the S phase checkpoint modulates multiple biological processes, including the repression of late-firing replication origins, cell cycle progression, the production of deoxynucleotide triphosphates (dNTPs), the transcription of DNA damage response genes, and inhibition of homologous recombination. Accordingly, activation of Rad53 triggers pseudohyphal growth, since in *S. cerevisiae* or *C. albicans* mutants deficient in this kinase showed a drastic reduction of filamentation under genotoxic stress conditions (19, 20).

Here, we demonstrate that many, but not all, clinical isolates of *C. auris* are capable of pseudohyphal growth when treated with genotoxins such as HU, MMS, or the clinically relevant fungistatic 5-fluorocytosine (5-FC). Deletion mutants of genes involved in the S phase checkpoint, *RAD9* and *MRC1*, or homologous recombination, *RAD51* and *RAD57*, allowed us to probe whether a functional S phase checkpoint is required for filamentous growth in *C. auris*. Our work provides the first insight into how genome stability maintenance supports cell growth and proliferation, and what triggers morphogenetic switching in the newly emerged fungal pathogen of humans *C. auris*.

## RESULTS

### *C. auris* produces pseudohyphae under genotoxic stress

Because the ability to switch between unicellular and filamentous forms plays a role in pathogenesis in some fungi (reviewed in (11, 12)), we were interested whether *C. auris* has the capability to form filaments as well. Several conditions, which induce hyphal growth in *C. albicans*, such as incubation at 37 °C, and Lee’s medium at pH3.5 or pH6.5, as well as media containing serum, isoamyl alcohol, or bleocin, were tested, but none of these triggered filamentous growth in the S. Asian (clade I) *C. auris* strain UACa11 (data not shown) (15). Likewise, growth at 25 °C did not produce filaments as described previously for a *C. auris* clinical isolate (21). However, using strain UACa11, we observed filamentous growth in the presence of sublethal concentrations of genotoxic drugs affecting DNA replication progression or inducing DNA damage (HU, MMS, and 5-FC) (Fig. 1). HU inhibits the activity of ribonucleotide reductase (22), and thus induces a depletion of the dNTP pools (23). MMS creates bulky adducts by alkylating DNA that interfere with fork progression (24). 5-FC is converted into fluorouracil in the cell and perturbs RNA and DNA biosynthesis (25). The filaments observed in *C. auris* show characteristics attributed to pseudohyphae (9, 11). In contrast to true hyphae, filaments in *C. auris* are wider than the diameter of a yeast cell, do not present parallel sides, septa between neighbouring cells show visible indentations, and actin patches are not accumulating at the growing tip (Fig. 1). Moreover, nuclear divisions seem to occur at mother-daughter junctions, but this phenotype is often complicated by nuclear division defects due to the genotoxin treatments (Fig. 1).

**Figure 1.**
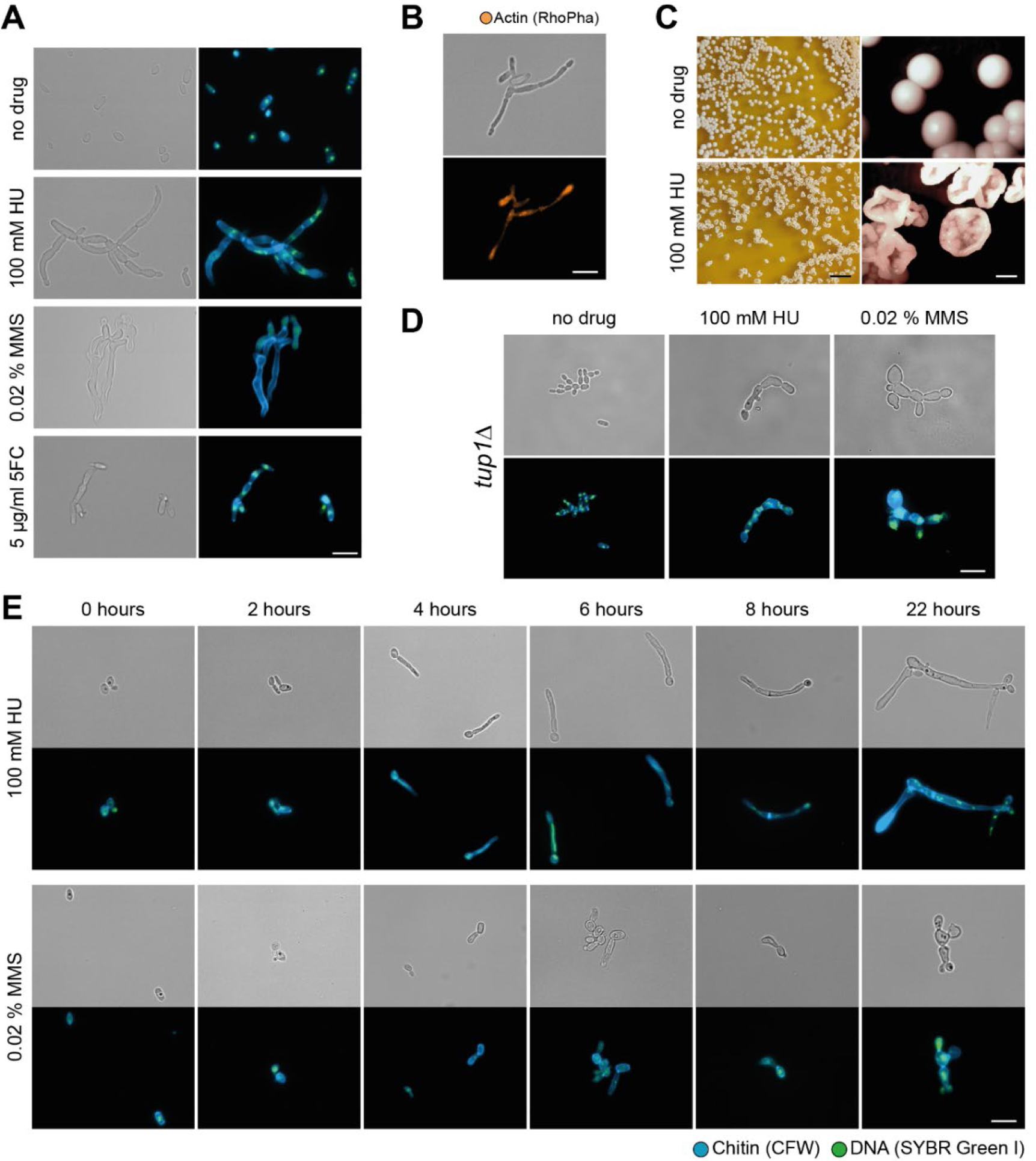
Filamentation of *Candida auris* (UACa11 and its mutant derivatives) in the presence of genotoxic drugs. (A) Microscopy images of *C. auris* filamentation after growing strain UACa11 on YPD plates with or without the addition of the indicated drug after 3 days at 30 °C; bright-field image on the left and merged fluorescent image (Chitin stained by calcofluor white, CFW, in blue, and DNA stained by SYBR Green I in green) on the right. (B) Cells grown on YPD plates containing 100 mM HU for 3 days at 30 °C stained for actin using rhodamine phalloidin (RhoPha) (bottom image), bright-field image on top. (C) Details of colonies from strain UACA11 grown on YPD plates in the absence and presence of 100 mM hydroxyurea (HU) for 6 days at 30 °C, scale bar on the left represents 10 mm, and scale bar on the right 1 mm. (D) *tup1*Δ cells treated and stain as wild-type cells in (A); bright-field images top row, fluorescent images bottom row. (E) Microscopy images of representative filaments of *C. auris* UACa11 formed in liquid culture; cells stained as in (A). After arrest in G1, cultures were grown for 165 min in YPD before indicated drugs were added (time-point 0 hours); bright-field images corresponding top rows, fluorescent images corresponding bottom rows. Scale bars in (A, B, D, E) represent 10 μm.

Within 6-8 h of growth in YPD containing 100 mM HU daughter cells started to show hyperpolarized growth, this results in almost 100 % filamentation after overnight culture (Fig. 1E), similar to *C. albicans* (18, 20). In this early filamentation phase generally one or two nuclei (SYBR Green-stained DNA signals) were observed within the same cell, occasionally very large high-intensity or stretched out DNA signals can be seen (Fig. 1E). After 22 hours constrictions at septa became evident throughout the filaments, new buds emerged from seemingly random locations along the filament; these cultures contained almost no separated (yeast) cells indicating a defect in cell division (Fig. 1E). Cells were often multi- or anucleate, suggestive of karyokinesis defects. A similar phenotype was generated by the addition of 0.02 % MMS (Fig. 1E), although the formation of filaments occurred more slowly and the resulting pseudohyphae tended to be shorter and more irregular than with HU treatment. Moreover, after 22 hours in MMS round and elongated yeast cells were still present in the cultures. Importantly, the number and length of filaments decreased over time when cells were grown on plates containing genotoxins (Fig. S1). On solid media filaments can also be observed without stress, as previously reported (26), but these are rare and could be explained by sporadic DNA damage. Strikingly, occasionally giant round cells were also observed in the presence of genotoxic drugs on solid media. In accordance with the cellular phenotype, the colony morphology after 5-6 days on solid medium is rougher when HU is present compared with the typical smooth colony appearance (Fig. 1C). This was not true in presence of MMS or 5-FC, either because these filaments were shorter, less abundant, or because they were more short-lived than the ones treated with HU (Fig. S1).

### *C. auris* lack key genes associated to hyphal growth

Filamentation is a complex mechanism, usually triggered by environmental conditions, involving hundreds of genes in *S. cerevisiae* and *C. albicans* (27–29). Among them, a group of well-studied key regulators known as hypha-specific genes (HSGs). Previously, Muñoz and co-workers, described that some HSGs essential for true hyphal growth, such as *ECE1* or *HWP1* are absent in the *C. auris* genome (30). We further explored the presence of key genes in filamentation in the *C. auris* genome (Table S1). Besides *ECE1* and *HWP1*, *FLO11*, *EED1*, and *HWP2* were also absent in *C. auris*. In *S. cerevisiae* Flo11 is a key factor for pseudohyphal growth in response to nutrient limitation (31, 32). Intriguingly, in *C. albicans*, which also lacks Flo11, Hwp2 seems to cover this function (33). *UME6*, which codes for a Zn(II)2Cys6 transcription factor involved in filamentation regulation, is present in *Candida* species capable of forming true hyphae (*C. albicans*, *C. tropicalis*, *C. dubliniensis*) and species unable to form true hyphae (*C. parapsilosis*, *C. orthopsilosis*, *C. lusitaniae*). Importantly, in the latter only the C-terminal Zn(II)2Cys6 domain is well conserved; this is also true for *C. auris* (Fig. S2).

Tup1 is a transcriptional repressor for filamentous growth in *C. albicans* acting on several HSGs, and *tup1*Δ strains grow constitutively as true hyphae (34, 35). In a *C. auris tup1*Δ strain no filaments were observed when grown without stress (Fig. 1D). However, strings of yeast cells were frequently observed, indicating that cells cannot separate properly. Under genotoxic stress, the *tup1*Δ mutant formed pseudohyphae similar to the parental strain. Altogether, this suggests that filamentation in *C. auris* is likely regulated differently than in *C. albicans*.

### *C. auris* has a functional S phase checkpoint

Genotoxic drugs induce replication fork perturbations and/or DNA damage, leading to cell cycle arrest by triggering the S phase checkpoint. Genes encoding S phase checkpoint factors – *MEC1* (XP_028890424), *RAD9* (XP_028889586), *MRC1* (XP_028891779), and *RAD53* (XP_028891118) – and homologous recombination proteins – *RAD51* (XP_028892133) and *RAD57* (KND99929) – were found in the *C. auris* genome (Figs. S3-4, Table S1). Null mutants of *RAD9*, *MRC1*, *RAD51*, and *RAD57* were obtained by deleting the open reading frames in the *C. auris* UACa11 strain (Fig. S5). Unfortunately, three attempts to generate *mec1*Δ or *rad53*Δ null mutants checking >300 transformants each were unsuccessful, possibly because those genes are essential in *C. auris*. The *rad51*Δ and *rad57*Δ strains showed very similar phenotypes (Fig. S6), for detailed analysis we focussed on the *rad51*Δ strain.

The mutant phenotypes were characterized using sublethal concentrations of various genotoxic drugs (Fig. 2). None of the mutants showed a conspicuous growth defect in the absence of genotoxins (Fig. 2). The *rad9*Δ strain showed sensitivity only to MMS and bleocin, indicating a role in responding to double-stranded DNA breaks (Fig. 2). In contrast, *mrc1*Δ displays sensitivity to drugs inducing replication-associated damage (HU, MMS, 5-FC) but grows like the parental control strain on bleocin (Fig. 2). Growth of *rad51*Δ was severely affected in presence of HU and MMS, but only moderately so in presence of bleocin, and was almost indistinguishable from wild type on 5-FC (Fig. 2). These mutant *C. auris* phenotypes are similar to the ones observed in the corresponding *S. cerevisiae* and *C. albicans* mutants (20, 36–39). However, Mrc1 seems to play a lesser role in *C. auris*, since in the other two yeasts *mrc1*Δ strains were more sensitive to MMS, and in *C. albicans* this mutant showed a growth defect even when genotoxic challenges were absent (20, 36–39). Taken together, these results suggest that the S phase checkpoint is conserved in *C. auris*, albeit with a somewhat reduced importance of Mrc1.

**Figure 2.**
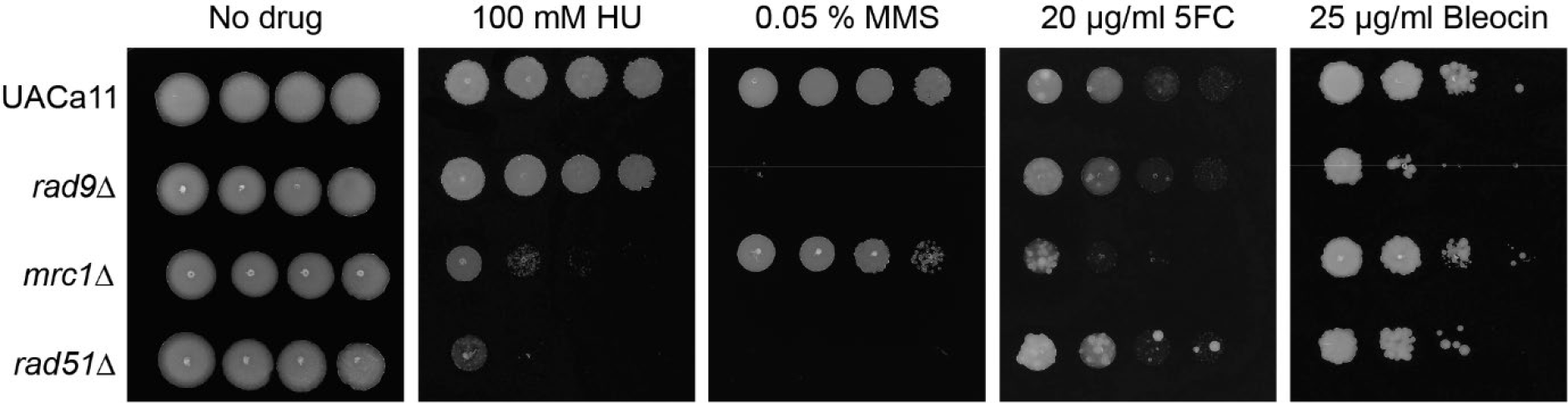
*Candida auris rad9*, *mrc1*, and *rad51* mutants show sensitivity to genotoxic drugs. Growth analysis by spot assays of wild type (UACa11) and the indicated deletion mutants in the presence and absence of genotoxic drugs. 10-fold serial dilution of *C. auris* cells were grown on YPD plates containing the indicated drug for 3 days at 30 °C.

The S phase checkpoint slows down the cell cycle in response to DNA damage or DNA replication inhibition. The role of *C. auris* Rad51, Rad9 and Mrc1 for cell cycle arrest in response to genotoxic stress was further studied. Wild-type *C. auris* (UACa11) cultures can be arrested in G1 by nitrogen starvation; almost 100 % of cells grown without a nitrogen source arrested in G1 after 7 hours, and remained in G1 during prolonged (24 hours) starvation (Fig. S7A). Upon return to favourable growth conditions (YPD), wild-type cultures had a lag phase of ~2.5-3 hours (Fig. S7B). Therefore, we grew wild-type and mutant yeast cultures for 165 min in YPD after G1 arrest before we started the experiments (time-point 0 hours) by adding 100 mM HU or 0.02 % MMS as genotoxic challenge; no drugs were added for the control. Cells were harvested at different time-points and the replication status of the cells was determined by flow cytometry (Fig. 3). In unperturbed conditions, wild type (UACa11), *rad9*Δ, and *rad51*Δ cells progressed from G1 to G2 within the first hour after re-entering the cell cycle (Fig. 3). This is in line with these mutants showing wild-type growth in solid medium in the absence of genotoxins (Fig. 2). In *mrc1*Δ a larger fraction of the cell population was in S phase, which suggests that S phase might last longer in this mutant similar to *S. cerevisiae* and *C. albicans* (20, 40, 41). Intriguingly, in *C. auris* this does not result in a notable growth defect on solid medium (Fig. 2).

**Figure 3.**
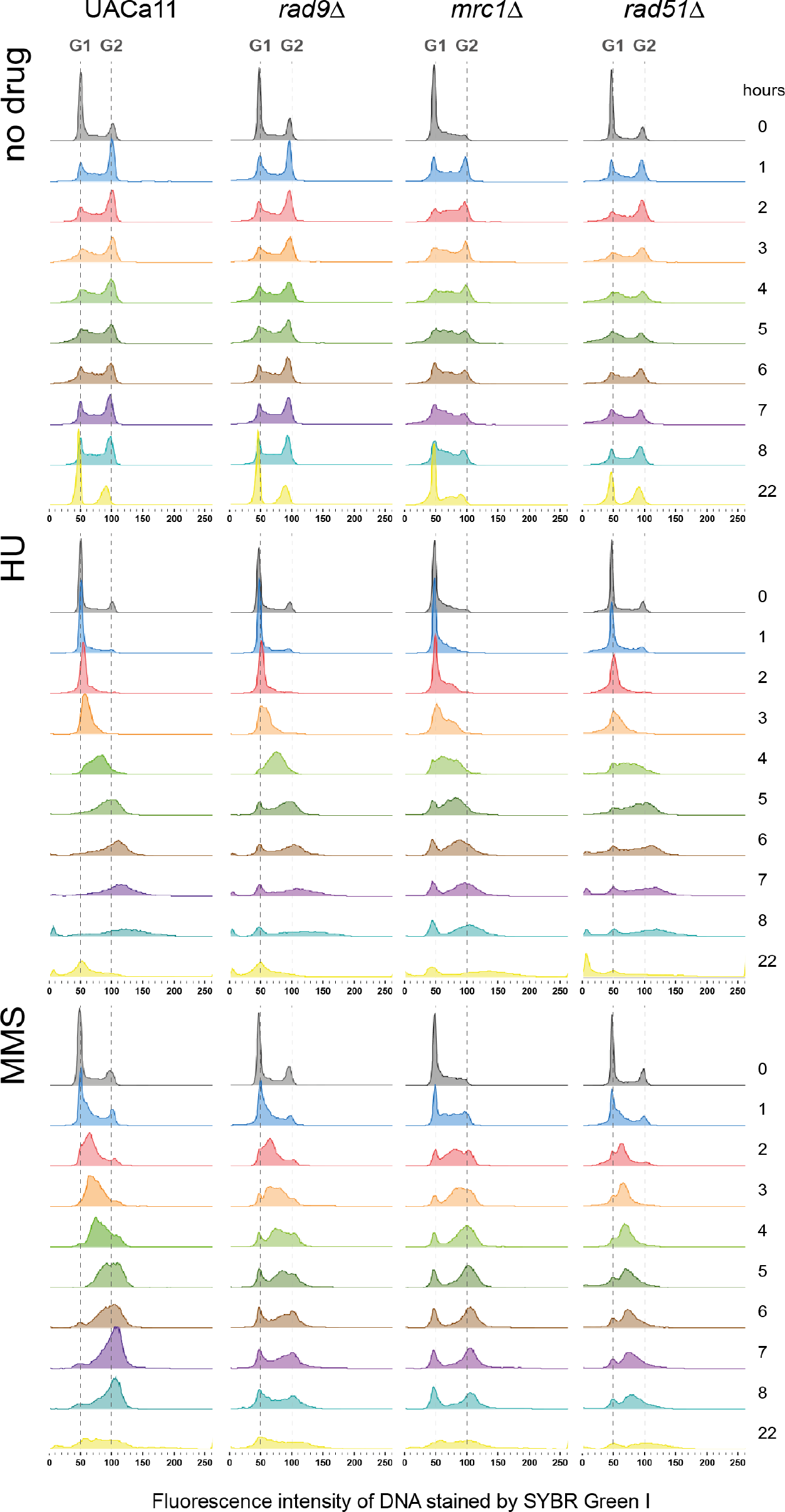
Cell cycle progression of *Candida auris* mutants under genotoxic stress. Histograms showing cell cycle profiles obtained by flow cytometry of wild type (UACa11) and the indicated mutant derivatives. Cells, previously arrested in G1 by nitrogen starvation, were transferred to fresh YPD broth and grown for 165 min at 30 °C to restart the cycle before adding 100 mM HU, 0.02 % MMS, or no drug (time-point 0 hours). Cells were harvested at indicated time-points and DNA stained using SYBR Green I. Amount of DNA is expressed as fluorescence intensity. Approximate positions of G1 and G2 peaks are indicated with dotted lines.

In the parental wild-type strain the S phase checkpoint is functional, and in the presence of HU and MMS cell cycle progression is slower than in unperturbed conditions, with a large fraction of cells in S phase between 4 and 6 hours (Fig. 3). Eventually, the checkpoint adapts, and cells move to G2 (Fig. 3). This is also largely true for *rad9*Δ and *rad51*Δ in the presence of HU, indicating a minor role in the HU response (Fig. 3). The *mrc1* null mutant behaves differently and shows a slower progression than the parental strain, since after 6 hours the majority of cells are still in S phase (Fig. 3). In the *rad51*Δ mutant most of the cells were arrested in S phase under MMS-treatment, whereas cell cycle progression in the *rad9*Δ and *mrc1*Δ mutants were similar to the parental strain under these conditions (Fig. 3). However, in all the mutants, but not the wild type, a small population of G1 cells was always present during genotoxin treatments (Fig. 3). This could mean that some cells never enter the cell cycle after the G1 arrest or escape the S phase checkpoint early-on and progress through the cell cycle to G1 quickly; these possibilities are not mutually exclusive. Indications that the latter possibility actually occurs, are (I) that these G1 populations grow over time in *rad9*Δ and *mrc1*Δ mutants under both genotoxic conditions, and (II) that it also occurs in wild type when challenged with MMS. This indicates that *rad9*Δ and *mrc1*Δ are indeed defective in the S phase checkpoint. At 22 hours the cycle seems to be partially restored (Fig. 3), though the presence of filaments and increased cell death after genotoxic treatment confounds the interpretation of this result. Taken together, these results support that a functional S phase checkpoint exists in *C. auris*.

### The S phase checkpoint is involved in pseudohyphal growth

Studies in *C. albicans* and *S. cerevisiae* have shown that HU and MMS treatment induces pseudohyphal growth dependent on activation of the S phase checkpoint, because a reduction of filamentation was described for strains carrying mutations in the gene coding for the S phase checkpoint kinase Rad53 (19, 20). The ability of mutant *C. auris* to produce filaments was tested in liquid (Fig. 4A) and on solid media (Figs. 4B, S1). As described above, wild-type cells of strain UACa11 form pseudohyphal filaments upon treatment with various genotoxins (Fig. 1).

**Figure 4.**
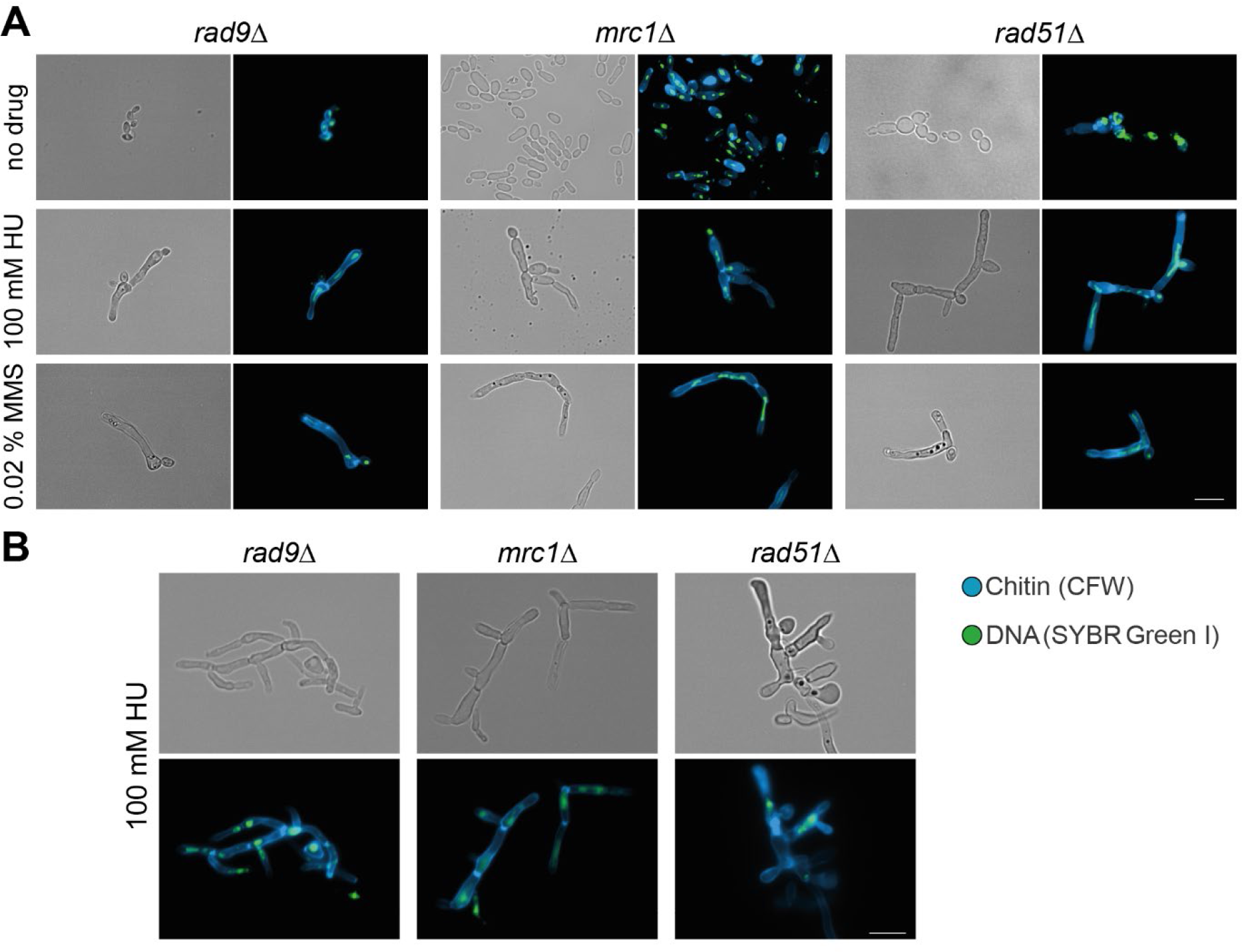
Microscopic analysis of filament formation in *Candida auris rad9*, *mrc1*, and *rad51* mutants (UACa11 background). (A) Representative microscopy images of *C. auris* filaments after growing *rad9*Δ, *mrc1*Δ, and *rad51*Δ cells in YPD broth with or without the addition of the indicated drug for 18-20 hours at 30 °C; bright-field images in the left columns, and merged fluorescent images (Chitin stained by calcofluor white, CFW, in blue, and DNA stained by SYBR Green I in green) in the right columns. (B) Representative microscopy images of *C. auris* filaments after growing *rad9*Δ, *mrc1*Δ, and *rad51*Δ cells on YPD plates containing 100 mM HU after 3 days at 30 °C; bright-field images on top, and merged fluorescent images (Chitin stained by calcofluor white, CFW, in blue, and DNA stained by SYBR Green I in green) at the bottom. Scale bars represent 10 μm.

*C. auris rad9*Δ cells grew as yeasts without genotoxic stress and as pseudohyphae in the presence of HU, similar to the wild type (Fig. 1) and to *C. albicans rad9*Δ (20). Intriguingly, *rad9*Δ cells produced filaments growing in liquid media containing MMS, as observed in the wild-type parent, but failed to form filaments on solid medium with MMS even after 2 days. That partially contrasts with the situation in *C. albicans*, where *rad9*Δ cells were somewhat compromised for filamentous growth in response to MMS compared to wild type, but would still form short pseudohyphae (20). After three days on solid medium containing 0.02 % MMS, a large fraction of *rad9*Δ cells became very large and round (giant cells), most of these giant cells are gone after seven days of culture (presumably because giant cells have a low viability). MMS induces the formation of giant cells in the wild type as well, although at much lower frequency, indicating that the lack of Rad9 enhances the production of this cell type.

*C. auris mrc1*Δ cells were elongated, but not fully filamented, after growth of 18-20 hours in liquid media without genotoxins, in comparison to the round or oval shape of wild-type cells. This differs somewhat from *C. albicans* where Mrc1-defective cells displayed pseudohyphal growth without stress (20). However, on solid media without genotoxic stress pseudohyphae were observed in *mrc1*Δ cells. In the presence of HU or MMS, the phenotype of *mrc1*Δ mutant was almost identical to the wild-type parent.

As observed in *rad52*Δ and *rad51*Δ cells in *C. albicans* (37, 42), a sizeable fraction of cells in the *C. auris rad51*Δ mutant showed constitutive pseudohyphal growth in unstressed conditions. The remainder of the cells showed a yeast phenotype, or were somewhat elongated often of aberrant shape (e.g. giant cells, very wide and elongated cells). When treated with HU or MMS, this mutant formed pseudohyphae to a similar extent as wild type, but the filaments tended to be longer compared to the parental UACa11 strain, especially on solid medium after 14 days (Fig. S1).

Overall, this suggests a role of the S phase checkpoint in *C. auris* filamentous growth, although the involvement of this pathway slightly differs from what has been observed in *C. albicans*.

### *C. auris* filamentation is strain-dependent

Strikingly, filamentation in *C. auris* was strain-dependent. We tested the filamentation of 22 different *C. auris* clinical isolates (Table S2) covering the four major clades (43) by treatment with HU (Figs. 1, 5, S8). Among them, different grades of filamentation were observed: some strains showed longer filaments (UACa11 or UACa24) than others (UACa6 or UACa23), some were straight and thin (UACa11 or UACa24), or wider and shorter growing as chains of (elongated) cells (“bubbles”) (UACa7 or UACa22), some formed aberrant cell shapes more frequently (UACa4 or UACa25). The clade III strain UACa10 seems to be unable to produce filaments when treated with genotoxins, it tended to produce bigger cells than in unperturbed conditions though. The grade of filamentation was not obviously correlated with a particular clade, since isolates from the same clade showed different phenotypes.

**Figure 5.**
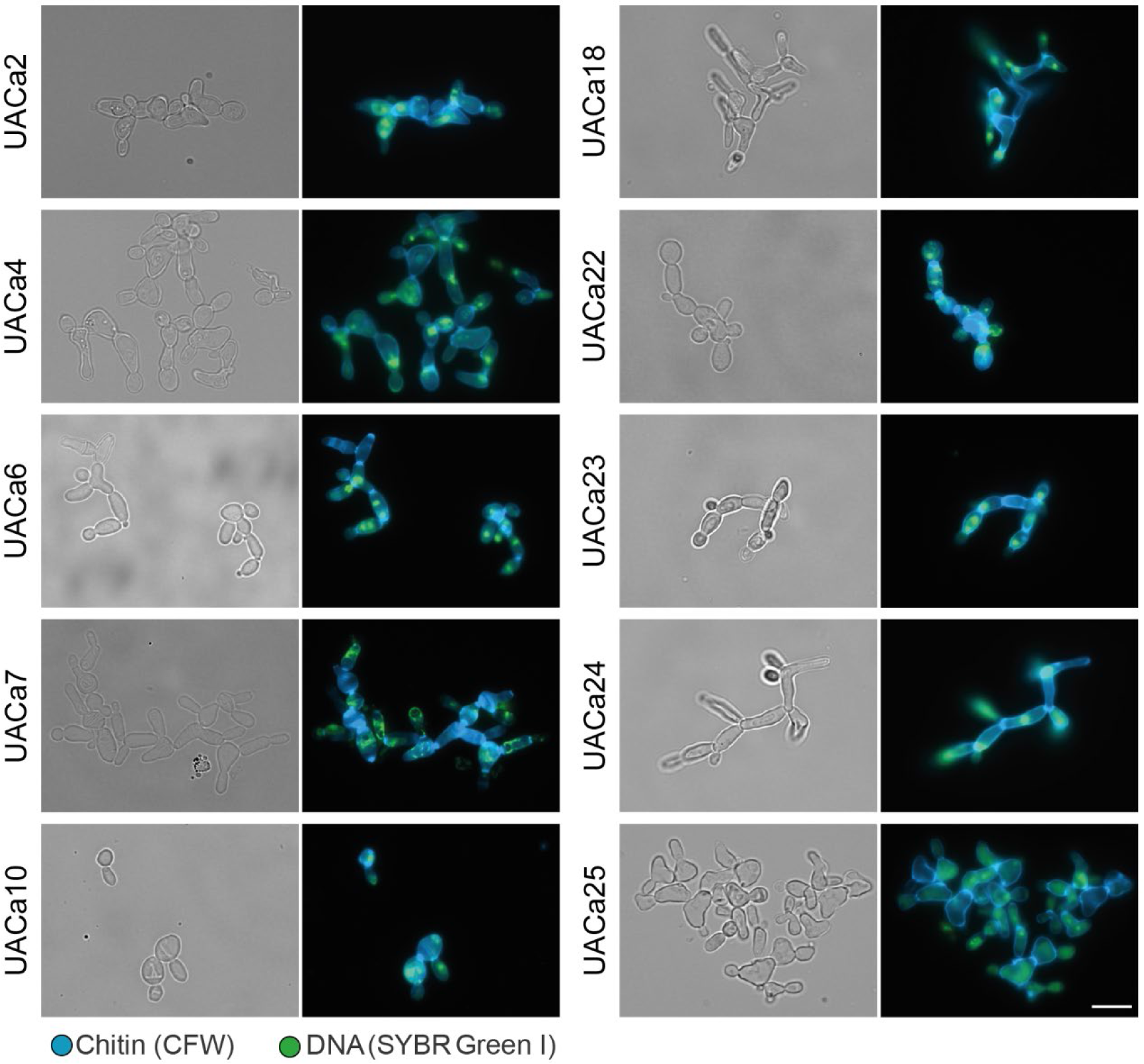
Different degrees of filamentation in *Candida auris* clinical isolates. Representative microscopy images of selected clinical isolates (see Table S2) after growth in YPD broth containing 100 mM HU for 18-20 hours at 30 °C; bright-field images in the left columns, and merged fluorescent images (Chitin stained by calcofluor white, CFW, in blue, and DNA stained by SYBR Green I in green) in the right columns. Scale bars represent 10 μm.

**Figure 6.**
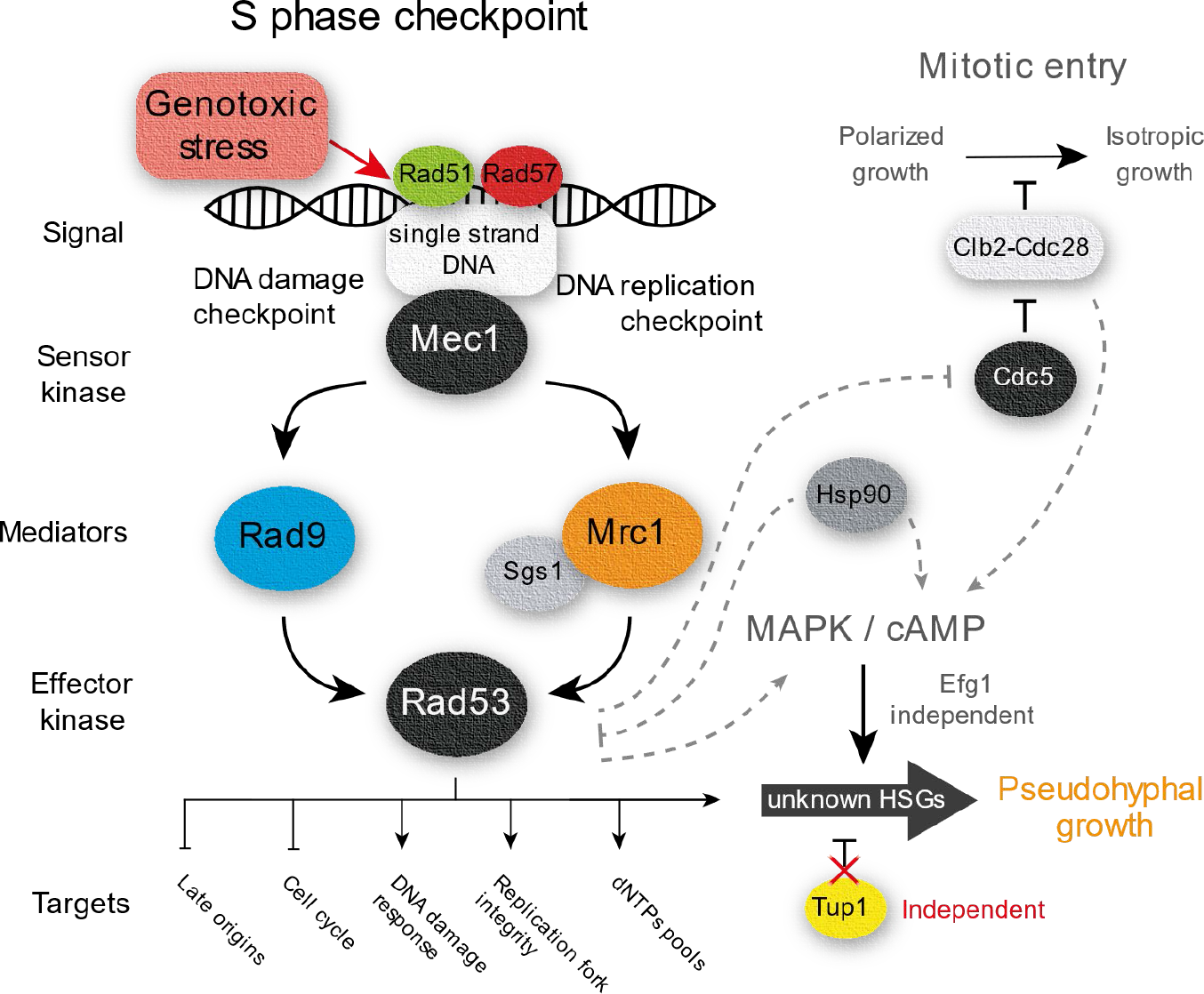
Schematic representation S phase checkpoint expected for *Candida auris*. Working model of the S phase checkpoint according to results in this study and related literature in yeast. Proteins tested in this study are in full colour. Possible S phase checkpoint downstream pathways related to filamentous growth are connected by dotted lines. HSGs (Hyphal-Specific Genes).

To evaluate if these strain-dependent filamentation phenotypes are due to differences in cell cycle progression under genotoxic stress, we selected five strains (UACa10, UACa11, UACa18, UACa22 and UACa25) representing all four clades and showing different grades of filamentation (Figs. 1, 4). As before, these strains were subjected to nitrogen starvation to arrest them in G1, and after pre-growth in liquid media for 165 min, genotoxins were added. In the presence of HU, the cell cycle in strains UACa10, UACa22 and UACa25 progressed ~1 hour faster than in strain UACa11 (Fig. S9). When treated with MMS the cell cycle progression in UACa25 was ~2 hours faster (Fig. S9). In the slow-growing clade II isolate UACa18 a fraction of the cell population remained in G1 after MMS treatment (Fig. S9), which indicates that the re-start of the cell cycle after G1 arrest might be slower in this strain. Although we observed some subtle differences between the *C. auris* isolates, these do not explain their substantial differences in filamentation.

Therefore, we looked for SNPs between two strains from the same clade showing different grades of filamentation, which might explain this differential cellular behaviour (isolates from different clades are too variable, and it would be too difficult to filter out the noise from these data) (43–45). We selected two strains from clade I (South Asia), UACa1 and UACa4, with different capabilities of forming pseudohyphae, which differ by 298 SNPs. Intriguingly, of a total of 99 non-synonymous SNPs within open reading frames we found 23 genes with a predicted/demonstrated role in filamentation and one hypothetical protein of unknown function with a SNP causing a premature stop codon in UACa1 (Table S3). Unfortunately, this makes it unfeasible to determine a single target gene as reason for the phenotypic differences between these two strains, and suggests that several pathways could be involved in enabling or preventing a particular strain to filament under stress conditions.

## DISCUSSION

Due to its recent emergence, there is a lack of understanding about the life cycle of *C. auris* which impedes a full comprehension of its origin, cellular behaviour, and pathogenic traits. Due to the evolutionary distance to the best-studied *Candida* species, *C. albicans* (30), inferences from research on *C. albicans* are not transferable to *C. auris*. This certainly is the case for the morphological switch between yeast cells and filaments. Most cues causing filamentation in *C. albicans* do not work in *C. auris* (15, this study). Accordingly, several genes essential for filamentation in *C. albicans* and *S. cerevisiae*, such as *EED1*, *FLO11*, *HWP1*, *HWP2* or *ECE1* (11, 12) are missing from the *C. auris* genome (Table S1) (30) and some important regulatory determinants of filamentation, such as Ume6, show conspicuous differences between *C. auris* and *C. albicans* (Fig. S2). Accordingly, *tup1*Δ cells do not trigger constitutive filamentation in *C. auris* (Fig. 1D) as it does in *C. albicans* (34). However, untreated *tup1*Δ cells, showed defective cell separation. This phenotype might be due to overexpression of *HGC1*, encoding for a G1 cyclin, which controls the expression of septum-degrading enzymes in *C. albicans* (46) and its expression is kept off by the repressor complex Tup1/Nrg1 (47). Altogether, these results indicate that *C. auris* is incapable of inducing filamentous growth under the same stimuli as *C. albicans*.

However, we observed that genotoxic stress triggers filamentation in most *C. auris* isolates tested (Figs. 1, 5, S8). These filaments displayed characteristics attributed to pseudohyphae (reviewed in (9, 11)) as previously described in presence of HU (48). Indeed, the presence of genotoxic drugs, such as MMS or HU, also trigger pseudohyphal growth in *C. albicans* and *S. cerevisiae* (18–20, 49, 50).

In fungi, the switch from yeast to filamentous growth is signalled through the mitogen-activated protein kinase (MAPK) and the fungal cyclic AMP (cAMP)-protein kinase A (PKA) pathways. Additionally, other pathways, such as the sucrose-nonfermentable (SNF), TOR, Hog1, and Rim101 pH-sensing pathways, influence filamentation (reviewed in (11–13)). This process involves ~700 genes for pseudohyphae formation In *S. cerevisiae* (27, 28), and more than 2,000 in *C. albicans* filamentous growth (29). However, mutation of key genes regulating filamentous growth, such as *HGC1*, *UME6*, *FLO8*, *TEC1*, *NRG1*, *CPH1* (*STE12* in *S. cerevisiae*) or *EFG1* do not affect HU-induced pseudohyphae formation (19, 49, 50). Therefore, pseudohyphal growth upon genotoxic stress apparently involves, at least partially, different mechanisms.

Genotoxic drugs can induce a variety of DNA damage types and/or perturbation of DNA replication forks, thus triggering the S phase checkpoint. As part of the checkpoint response the kinase Rad53 is activated, which leads to a temporal cell cycle delay until the issue is resolved and the cell cycle can continue (reviewed in (17)). A delay of the cell cycle in S phase and a subsequent transient arrest in G2/M were observed in various *C. auris* strains in the presence of the genotoxic agents HU and MMS (Figs. 3, S7); this suggests that *C. auris* has a functional S phase checkpoint. Furthermore, mutation of the S phase checkpoint genes *MRC1* and *RAD9* leads to a defective cell cycle arrest, since some cells seem to progress quickly to G1 under genotoxic stress (Fig. 3) similar to the same *C. albicans* or *S. cerevisiae* mutants (20, 38, 40, 41, 51). Although activation of Rad53 by DNA damage or perturbed DNA replication forks is sensed differently, both types of DNA lesions share the molecular signal that triggers the response: an accumulation of single-strands DNA (ssDNA) which acts as signal for recruiting and later activating Mec1 (52–54). Homologous recombination is required to repair the DNA lesions generated under genotoxic stress and, therefore, Rad51 and Rad52 are required to prevent an excess of ssDNA (52, 55, 56). Indeed, in *C. auris rad51*Δ cells were arrested in S phase during MMS treatment, which suggests an inability to restore the DNA lesions and, therefore, a constitutive activation of Rad53. Moreover, similar to *rad9*Δ or *mrc1*Δ strains and in contrast to wild type, a fraction of the genotoxin-stressed *rad51*Δ cell populations seem unable to re-start the cell cycle after the G1 arrest or move through the cell cycle without delay (see above) (Fig. 3). Altogether, our results suggest that similar to other ascomycetes, *C. auris* has fully functional S phase checkpoint and homologous recombination pathways.

Pseudohyphal growth in response to genotoxins is S phase checkpoint-dependent, as *rad53* mutants in *C. albicans* or *S. cerevisiae*, and *mec1*Δ in *S. cerevisiae* showed a drastic decrease of filamentation under genotoxin treatment (19, 20). Likewise, strains defective for *RAD9* or *MRC1* also showed alterations of morphology in the presence of genotoxic stress in *C. albicans* (20). After MMS treatment a *C. albicans RAD9*-defective mutant forms filaments which are considerably shorter than in wild type. In contrast, filamentation of a *C. auris rad9*Δ mutant was not different from wild type after 24 h of MMS-treatment in liquid culture (Fig. 4). However, on plates fewer pseudohyphae were observed in the *C. auris rad9*Δ mutant in comparison to wild type after long-term exposure to MMS (>2 days), albeit a higher proportion of yeast cells become enlarged in the mutant (Fig. S1). This cell enlargement was obvious in, but not restricted to, *rad9*Δ and could occasionally be observed in wild type in response to MMS applied for periods longer than 2 days (Fig. S1). In other fungi, the presence of enlarged round cells has been described as Titan cells in *Cryptococcus neoformans* (57, 58), or Goliath cells in *C. albicans* (59). Further studies will be necessary to elucidate, whether these large cells in *C. auris* resemble Titan or Goliath cells or are something completely different. Taken together this would suggest that in *C. albicans* Rad9 might control different or additional HSGs compared to the situation in *C. auris* (see below). The *C. auris mrc1*Δ mutant induced, at least partially, pseudohyphal growth in the absence of any genotoxic stress, otherwise it was indiscernible from wild type (Figs. 4, S1), this result is similar to observations in *C. albicans* (20, 60). Interestingly, a *C. albicans SGS1* mutant strain, which fails to activate Rad53 via Mrc1, forms filaments in unperturbed conditions (61). As a possible explanation, formation of DNA double strand breaks in *mrc1*Δ strains, and the subsequent activation of Rad53 through Rad9 has been suggested (40, 62). An S phase-independent role has also been described for Mrc1 though, regulating the replication initiation through interaction with Cdc7, a conserved kinase that triggers firing at each replication origin (63), which regulates the mitotic exit through interaction with Cdc5 (64). This could explain the filamentation observed in the *mrc1*Δ mutant (see below), and the larger S phase population observed in this mutant in unperturbed conditions (Fig. 3). Mutation of homologous recombination genes such as *RAD51*, *RAD52*, *MRE11*, or *RAD50*, causes constitutive pseudohyphae formation in *C. albicans* (37, 42, 61). We observed that deletion of *RAD51* and *RAD57* also triggered constitutive pseudohyphal growth in *C. auris* (Figs. 4, S6).

The S phase checkpoint is a complex process including several back-up mechanisms which could explain why *MRC1* and *RAD9* mutants are still able to arrest the cell cycle and produce filaments under genotoxic stress. Indeed, it has been reported that Mrc1 depends on Rad9 to stay activated for a long period, and that Rad53 is rapidly but only transiently activated by Mrc1 in *rad9*Δ cells, and is slowly, but continuously, activated by Rad9 in absence of *MRC1* (17, 51). Accordingly, the double mutant *mrc1*Δ *rad9*Δ is inviable (40, 62). That would explain our observations, under MMS-treatment, that in the short term *C. auris rad9*Δ produces filaments, due to activation of Rad53 by Mrc1, but in the long term, Mrc1 is not able to maintain this activation and cells lose their filamentation phenotype.

Mechanisms involved in pseudohyphal growth in response to S phase checkpoint activation are not well understood and further studies will be necessary. However, one mechanism could involve the constitutive activation of the Clb2-Cdc28 complex by Rad53 in response to genotoxic stress, through the polo kinase Cdc5 (65–67). The activation of Clb2-Cdc28 prevents the entry into mitosis and the associated switch from polarized to isotropic growth (68, 69), therefore, cells would be stuck in the apical growth phase, thus forming filaments. Hence, mutation of *CLB2*, *CDC28*, and *CDC5* triggers constitutive pseudohyphal growth in yeasts (49, 70, 71). Furthermore, the cAMP and MAPK pathways have been implicated in pseudohyphal growth in response to genotoxic stress via downstream regulators (18, 42, 50, 72). A plausible model for *C. auris* is depicted in Fig. 6; our model also takes into account that, in *C. auris*, constitutive pseudohyphal growth is triggered by downregulation of *HSP90*, a heat-shock family protein, which acts as chaperone and influences a diverse range of signal transducers (48). Inhibition of *HSP90* induces pseudohyphal growth via cAMP-PKA signalling in an Efg1-independent way in *C. albicans* (73) and, interestingly, a direct inhibition of Rad53 by Hsp90 has been observed in *S. cerevisiae* (74).

During infection, cells may encounter various conditions that lead to cell cycle arrest, either produced by the host or by other microorganisms cohabiting a given niche. Switching to filamentous growth might be advantageous, allowing cells to escape from such perturbations. However, to our knowledge, filamentous growth in the context of pathogenesis and colonization has only been described once in *C. auris* (21). This strain recovered from an infected mouse displayed filamentous growth at temperatures below 25 °C. However, none of our strains have developed filamentation under these conditions. Anyway, there currently is not enough data available for *C. auris* infection to fully appreciate the potential role of morphogenetic switching, which contributes to full virulence in *C. albicans* (reviewed in (11, 13, 14)). The different capacities of different strains to form filaments (21, this study) and the plasticity of the *C. auris* genome (43, 75) are indications of the flexibility and adaptability of this fungus suggesting that different clinical isolates could use morphogenetic switching during different phases of pathogenesis.

## METHODS

### Yeast strains and culture conditions

*Candida* strains used in this study are listed in Table S2. Yeast cells were grown at 30 °C on YPD plates (1 % yeast extract, 2 % mycological peptone, 2 % glucose, 2 % agar; Oxoid, Basingstoke, UK) or shaking at 200 rpm in YPD broth (same as plates, but without agar). Cell concentrations were determined by measuring optical density of the culture at a wavelength of 600 nm (OD_600_) on an Ultraspec 2000 (Pharmacia Biotech, Uppsala, Sweden) spectrometer (calibration defined OD_600_ = 1 to contain ~3 × 10^7^ *C. auris* cells/ml).

Spot assays were carried out on YPD agar adding the indicated drug where necessary. Cell concentrations were determined by OD_600_ after overnight growth in YPD broth, four 1:10 serial dilutions containing 10^2^ to 10^5^ cells were plated and grown at 30 °C for 3 days (unless indicated otherwise).

For filament visualization, ~1 × 10^7^ cells were grown for 18-20 hours in YPD broth, unless indicated otherwise, adding 100 μM HU, 0.02 % MMS, 5 μg/ml 5-FC. Other potential filamentation conditions tested were: Lee’s medium pH3.5 and pH6.5 (76); YPD (see above) or RPMI-1640 containing 20 % fetal calf serum (Sigma-Aldrich Corp., St. Louis, MO, USA); in YPD medium at temperatures of 25, 30 or 37 °C; presence of 0.5 % isoamyl alcohol in YPD; and 25 μg/ml bleocin in YPD. To check filamentation on solid media, fresh cells were streaked on YPD plates containing 100 μM HU, 0.02 % MMS or 5 μg/ml 5-FC and grown at 30 °C for times indicated.

### Cell cycle analysis and microscopy

To synchronize in G1, cells were grown in YPD broth overnight at 30 °C and 200 rpm. Then 1 × 10^7^ cells were washed with sterile water and transferred to nitrogen starvation conditions [6.7 g/L YNB without amino acids and without ammonium sulfate (Sigma-Aldrich) and 20 g/l glucose] and grown at 30 °C and 200 rpm. Samples were harvested by centrifugation (1,000 ×*g*, 2 min), and fixed adding 70% ethanol at the indicated time-points. Cells were stored at +4 °C before proceeding to flow cytometry (see below).

The cell cycle of cells previously arrested in G1 from different *C. auris* strains in the presence of genotoxic drugs were determined. For this purpose, 1 × 10^7^ cells, obtained after growing in YPD broth overnight at 30 °C and 200 rpm, were harvested, washed with sterile water, and transferred to nitrogen starvation conditions (see above) for 15-17 hours at 30 °C and 200 rpm. Then, 1 × 10^8^ cells were transferred to 10 ml of fresh YPD broth. To restart the cell cycle, cells were grown for 165 min at 30 °C and 200 rpm, after which 100 mM HU, 0.02% MMS were added (a third culture without any additions served as control). Cells were harvested by centrifugation (1000 ×*g*, 2 min), and fixed adding 70% ethanol, every hour from 0 to 8 h, and at a final time-point of 22 h. Cells were stored at +4 °C before proceeding to flow cytometry (see below). For flow cytometry yeast samples were prepared as previously described (75, 77).

For microscopy, cells were fixed by adding 1 ml 70% EtOH, and incubated for at least two hours at 4 °C. Fixed cells were pelleted (2 min, 1,000 ×*g*), and resuspended in sodium citrate buffer (14.7 g/l sodium citrate, pH 7.5). Cells were then treated with RNase A (250 μg per 1 × 10^7^ cells) and proteinase K (1,000 μg per 1 × 10^7^ cells) one hour at 50 °C, subsequently Triton-X 100 (Sigma-Aldrich) was added to a final concentration of 0.25%. Cells were stained for DNA (SYBR Green I, 1:5000; Sigma-Aldrich), and for chitin (1 μg/ml Calcofluor white; Sigma) at +4 °C overnight. For time course experiments, the same samples were used for flow cytometry and microscopy (adding 1 μg/ml Calcofluor white for microscopy). Rhodamine-Phalloidin staining was carried out as previously described (78) with a few modifications. Approximately 1 × 10^7^ cells from an overnight culture were fixed for 1 hour in 4 % formaldehyde (PBS-buffered) at room temperature, harvested by centrifugation (2 min, 1,000 *×g*) and resuspended in PBS before adding 10 μl Rhodamine Phalloidin (6.6 μM in Methanol) and 1 μl of DAPI (10 μg/ml). Samples were incubated in the dark for 1 hour, washed in PBS (2 min, 1,000 *×g*) twice. Finally, cells were resuspended in ProLong™ Diamond Antifade (Thermo Fisher Scientific, Waltham, MA, USA). Fluorescent images were taken on a Zeiss Axio Imager M2 microscope with a Zeiss 503 camera, and analyzed using ZEN 2 blue edition software (Carl Zeiss Microscopy, Jena, Germany). Standard brightfield microscopy was performed on a BX50 microscope (Olympus, Tokyo, Japan) equipped with an Infinity 1 camera (Lumenera, Ottawa, Canada), and analyzed using the Infinity capture software V 6.5.7 (Lumenera, Ottawa, Canada). Colony morphology was imaged on a Zeiss Stemi 2000-c binocular (Carl Zeiss Microscopy, Jena, Germany) equipped with an Infinity 1 camera (Lumenera, Ottawa, Canada).

### Nucleic acid manipulations

For fast gDNA extraction, cells from a single colony were suspended in 20 mM NaOH, boiled at 100 °C for 10 minutes, and incubated on ice for 10 minutes. Samples were then centrifuged (2 min, 1,000 ×*g*), and a small amount of supernatant was used for PCRs.

For high-quality gDNA extractions, cells from overnight cultures were harvested (2 min, 1,000 ×*g*), washed with double-distilled water (ddH_2_O). After removing the supernatant, acid-washed glass beads (Sigma-Aldrich) and DNA extraction buffer [1 M NaCl, 2% Triton x-100, 1 M Tris-HCl pH8, 0.5 M EDTA, 1 % SDS] were added. Cells were disrupted in a Vibrax VXR (IKA GmbH & Co. KG, Staufen, Germany) multi-tube mixer for 3 min before adding one volume of phenol:chloroform:isoamyl alcohol (25:24:1) (Sigma-Aldrich). Samples were mixed vigorously, and centrifuged (10 min, 12,000 ×*g*). The supernatant was transferred to a fresh reaction tube, and 0.1 volumes of 3 M sodium acetate pH 5.2 and 2 volumes of ice-cold 100% ethanol were added. After precipitation (30-60 min at –20 °C), samples were centrifuged (5 min, 12,000 ×*g*), and washed with 70% ethanol. Pellet were air-dried, resuspended in ddH_2_O, and 5 μl RNAse (10 mg/ml) were added followed by an incubation of 30-60 min at 37 °C. A small amount of a 1:10 dilution was used for subsequent PCRs.

Standard PCRs were carried out using DreamTaq PCR Master Mix (Thermo Fisher Scientific) following the manufacturer instructions. All primers used in this study (Table S4) were designed using the reference genome (https://www.ncbi.nlm.nih.gov/genome/?term=txid498019) and synthesized by Sigma-Aldrich.

### Gene deletion constructs and transformation

All gene deletion constructs were amplified using Phusion High-Fidelity PCR Master Mix (Thermo Fisher Scientific) as recommended by the supplier.

Genes were deleted using the nourseothricin-resistance marker *CaNAT1* flanked by 1.5-2 kb regions of homology to up- and downstream sequences of the target gene. *CaNAT1*, including *TEF*-promoter and -terminator sequences, was PCR-amplified from pV1025 (79) using oligonucleotides oUA315 and oUA316 (Table S4) and cloned into a *Bam*HI-linearized pFA6a backbone by NEBuilder DNA Assembly Master Mix (New England BioLabs, Ipswich, MA, USA). The assembly mix was transformed into NEB 10-beta *E. coli* cells (New England BioLabs) following the manufacturer instructions. The resulting plasmid, pALo218, was then used to amplify *CaNAT1* with oligonucleotides oUA353 and oUA354 (Table S4). Up- and downstream homologous regions of the target gene were PCR-amplified from gDNA of UACa11 (Table S2) using specific primers for each target gene (Table S4). The oligonucleotide primers on the 3’ end of the upstream region and at the 5’ end of the downstream region include short homologies to the *CaNAT1*-marker to assemble transformation cassettes via fusion PCR (80). The three PCR products (*CaNAT1*, up- and downstream homology regions) were combined in roughly equimolar amounts (~ 0.15 pmols) together with Phusion High-Fidelity PCR Master Mix (Thermo Fisher Scientific) containing 2.5 % DMSO and subjected to 8 PCR cycles (10 sec 98 °C, 30 sec 55 °C, 60 sec 72 °C). Nested primers (Table S4) under standard PCR conditions were used to amplify the final transformation cassettes.

Gene deletion constructs were transformed into *C. auris* strain UACa11 using a protocol developed for *C. albicans* (81) with small modifications. Briefly, cells from a culture grown in YPD to stationary phase were diluted 1:100 into fresh YPD broth, and incubated shaking at 30 °C until reaching mid-exponential phase (OD_600_ of 0.5-0.8). Cells were centrifuged (2 min, 1,000 *×g*), washed once with ddH_2_O, and resuspended in 100 mM lithium acetate. Transformation mixtures contained ~1 μg DNA of the transformation cassette, 100 μg carrier DNA (herring sperm DNA solution, Thermo Fisher Scientific), 37 % PEG (polyethyleneglycol, molecular weight 3,350), 100 mM lithium acetate, and ~1.5 × 10^8^ *C. auris* cells. The transformation mixture was incubated overnight (16-20 hours) at 30 °C. After a heat shock of 15 min at 44 °C, cells were pelleted by centrifugation (2 min, 1,500 *×g*), resuspended in YPD, and incubated shaking for 4 hours at 30 °C. Cells were then plated onto selective YPD agar containing 100 μg/ml nourseothricin (clonNAT; WERNER BioAgents GmbH, Jena, Germany) and incubated at 30 °C until transformants appeared. Gene deletion and correct integration were confirmed by PCR using primers located within the ORF of the target gene, and primers located within and outwith the transformation cassette (Fig. S5, Table S4).

### *In silico* analysis

Gene and protein sequences were obtained from the *C. auris* reference genome (assembly Cand_auris_B11221_V1) at the NCBI database (https://www.ncbi.nlm.nih.gov/genome/?term=txid498019[orgn]). Interestingly, *RAD57* was not present in this reference genome, but could be found int the draft genome of strain Ci6684 (82). Protein sequences of *Candida* spp were obtained from the *Candida* Genome Database (http://www.candidagenome.org/), or from Uniprot (https://www.uniprot.org/), and the *Saccharomyces* Genome Database (https://www.yeastgenome.org/) served as source for *S. cerevisiae* protein sequences. Homology scores of *C. auris* proteins against *C. albicans* and *S. cerevisiae* proteins (E-value, length aligned, and identities are shown in Table S1) were generated using the BLAST tools of the *Candida* or the *Saccharomyces* Genome Databases. Alignments were carried out by the MSAprobs method (default settings) using Jalview 2.10.5 (83). For SNP calling, Illumina sequencing reads from two clade I strains (UACa1, UACa4) were aligned (ZKR, NARG & AL; unpublished results) using BWA-MEM v0.7.12 (84) and processed with Samtools v0.1.19 view, sort, rmdup and index (85). SNPs were then detected using Pilon v1.22 (86) filtering the resulting variant call format (VCF) file for genotype 1/1 only. Low coverage (less than 10% of mean coverage), ambiguous positions and deletions were removed. The reference genome was annotated with Augustus v3.3.1 (87) *ab initio* gene prediction software and VCF annotator (http://vcfannotator.sourceforge.net) was used to predict the effect of the SNPs called on the annotated genes.

## ACKNOWLEDGEMENTS

We are grateful to Arunaloke Chakrabarti, Anuradha Chowdhary, Elizabeth Johnson (PHE), Takashi Kubota, and Shawn Lockhart (CDC) for providing strains. We thank Fei Long for skillful technical assistance. Flow cytometry was performed at the Iain Fraser Cytometry Centre (IFCC), University of Aberdeen (Raif Yuecel). Microscopy was done at the Microscopy & Histology Facility, University of Aberdeen (Kevin S. Mackenzie). This work was supported by a Wellcome Trust Seed Award to AL [grant number 212524/Z/18/Z], and the Medical Research Council (MRC) Centre for Medical Mycology at the University of Exeter [grant numbers MR/P501955/1, MR/N006364/1].

## AUTHOR CONTRIBUTIONS

Conceived and designed the study (GBR, NARG, AL); collected the data (GBR, ZKR); performed the analysis (GBR, ZKR, AL); drafted the manuscript (GBR, AL); revised the manuscript (GBR, ZKR, NARG, AL).

## REFERENCES

1. Satoh K, Makimura K, Hasumi Y, Nishiyama Y, Uchida K, Yamaguchi H. 2009. *Candida auris* sp. nov., a novel ascomycetous yeast isolated from the external ear canal of an inpatient in a Japanese hospital. Microbiol Immunol 53:41–44.

2. Kwon YJ, Shin JH, Byun SA, Choi MJ, Won EJ, Lee D, Lee SY, Chun S, Lee JH, Choi HJ, Kee SJ, Kim SH, Shin MG. 2019. *Candida auris* clinical isolates from South Korea: Identification, antifungal susceptibility, and genotyping. J Clin Microbiol 57:e01624–18.

3. Rhodes J, Fisher MC. 2019. Global epidemiology of emerging *Candida auris*. Curr Opin Microbiol 52:84–89.

4. Lone SA, Ahmad A. 2019. *Candida auris*—the growing menace to global health. Mycoses 62:620–637.

5. Kean R, Ramage G. 2019. Combined antifungal resistance and biofilm tolerance: the global threat of Candida auris 4:e00458–19.

6. Forsberg K, Woodworth K, Walters M, Berkow EL, Jackson B, Chiller T, Vallabhaneni S. 2018. *Candida auris*: The recent emergence of a multidrug-resistant fungal pathogen. Med Mycol 57:1–12.

7. Jackson BR, Chow N, Forsberg K, Litvintseva AP, Lockhart SR, Welsh R, Vallabhaneni S, Chiller T. 2019. On the origins of a Species: What might explain the rise of *Candida auris*? J Fungi 5:1–17.

8. Casadevall A, Kontoyiannis DP, Robert V. 2019. On the emergence of *Candida auris*: Climate change, azoles, swamps, and birds. MBio 10:e01397–19.

9. Sudbery P, Gow N, Berman J. 2004. The distinct morphogenic states of *Candida albicans*. Trends Microbiol 12:317–324.

10. Berman J. 2006. Morphogenesis and cell cycle progression in *Candida albicans*. Curr Opin Microbiol 9:595–601.

11. Sudbery PE. 2011. Growth of *Candida albicans* hyphae. Nat Rev Microbiol 9:737–748.

12. Cullen PJ, Sprague GF. 2012. The regulation of filamentous growth in yeast. Genetics 190:23–49.

13. Noble SM, Gianetti BA, Witchley JN. 2017. *Candida albicans* cell-type switching and functional plasticity in the mammalian host. Nat Rev Microbiol 15:96–108.

14. Thompson DS, Carlisle PL, Kadosh D. 2011. Coevolution of morphology and virulence in *Candida Species*. Eukaryot Cell 10:1173–1182.

15. Wang X, Bing J, Zheng Q, Zhang F, Liu J, Yue H, Tao L, Du H, Wang Y, Wang H, Huang G. 2018. The first isolate of *Candida auris* in China: clinical and biological aspects. Emerg Microbes Infect 7:93:0–8.

16. Kron SJ, Gow NA. 1995. Budding yeast morphogenesis: signalling, cytoskeleton and cell cycle. Curr Opin Cell Biol 7:845–855.

17. Pardo B, Crabbé L, Pasero P. 2017. Signaling pathways of replication stress in yeast. FEMS Yeast Res 17:1–11.

18. Bachewich C, Mantel A, Whiteway M. 2005. Cell cycle arrest during S or M phase generates polarized growth via distinct signals in *Candida albicans*. Mol Microbiol 57:942–959.

19. Jiang YW, Kang CM. 2003. Induction of *S. cerevisiae* filamentous differentiation by slowed DNA synthesis involves Mec1, Rad53 and Swe1 checkpoint proteins. Mol Biol Cell 14:5116–5124.

20. Shi Q-M, Wang Y-M, Zheng X De, Lee RTH, Wang Y. 2007. Critical role of DNA checkpoints in mediating genotoxic-stress–induced filamentous growth in *Candida albicans*. Mol Biol Cell 18:815–826.

21. Yue H, Bing J, Zheng Q, Zhang Y, Hu T, Du H, Wang H, Huang G. 2018. Filamentation in *Candida auris*, an emerging fungal pathogen of humans: passage through the mammalian body induces a heritable phenotypic switch. Emerg Microbes Infect 7:188.

22. Nyholm S, Thelander L, Gräslund A. 1993. Reduction and loss of the iron center in the reaction of the small subunit of Mouse ribonucleotide reductase with hydroxyurea. Biochemistry 32:11569–11574.

23. Poli J, Tsaponina O, Crabbé L, Keszthelyi A, Pantesco V, Chabes A, Lengronne A, Pasero P. 2012. dNTP pools determine fork progression and origin usage under replication stress. EMBO J 31:883–894.

24. Vázquez MV, Rojas V, Tercero JA. 2008. Multiple pathways cooperate to facilitate DNA replication fork progression through alkylated DNA. DNA Repair (Amst) 7:1693–1704.

25. Grem JL. 2000. 5-Fluorouracil: Forty-plus and still ticking. A review of its preclinical and clinical development. Invest New Drugs 18:299–313.

26. Sherry L, Ramage G, Kean R, Borman A, Johnson EM, Richardson MD, Rautemaa-Richardson R. 2017. Biofilm-forming capability of highly virulent, multidrug-resistant *Candida auris*. Emerg Infect Dis 23:0–3.

27. Jin R, Dobry CJ, McCown PJ, Kumar A. 2008. Large-scale analysis of yeast filamentous growth by systematic gene disruption and overexpression. Mol Biol Cell 19:284–196.

28. Ryan O, Shapiro RS, Kurat CF, Mayhew D, Baryshnikova A, Chin B, Lin ZY, Cox MJ, Vizeacoumar F, Cheung D, Bahr S, Tsui K, Tebbji F, Sellam A, Istel F, Schwarzmüller T, Reynolds TB, Kuchler K, Gifford DK, Whiteway M, Giaever G, Nislow C, Costanzo M, Gingras AC, Mitra RD, Andrews B, Fink GR, Cowen LE, Boone C. 2012. Global gene deletion analysis exploring yeast filamentous growth. Science (80-) 337:1352–1356.

29. Azadmanesh J, Gowen AM, Creger PE, Schafer ND, Blankenship JR. 2017. Filamentation involves two overlapping, but distinct, programs of filamentation in the pathogenic fungus *Candida albicans*. G3 Genes, Genomes, Genet 7:3797–3808.

30. Munoz JF, Gade L, Chow NA, Loparev VN, Juieng P, Farrer RA, Litvintseva AP, Cuomo CA. 2018. Genomic basis of multidrug-resistance, mating, and virulence in *Candida auris* and related emerging species. Nat Commun 9:5346.

31. Lo W, Dranginis AM. 1998. The cell surface flocculin Flo11 is required for pseudohyphae formation and invasion by *Saccharomyces cerevisiae*. Mol Biol Cell 9:161–171.

32. Vinod PK, Sengupta N, Bhat PJ, Venkatesh K V. 2008. Integration of global signaling pathways, cAMP-PKA, MAPK and TOR in the regulation of *FLO11*. PLoS One 3:e1663.

33. Younes SS, Khalaf RA. 2013. The *Candida albicans* Hwp2p can complement the lack of filamentation of a *Saccharomyces cerevisiae flo11* null strain. Microbiol (United Kingdom) 159:1160–1164.

34. Braun BR, Johnson AD. 1997. Control of filament formation in *Candida albicans* by the transcriptional repressor TUP1. Science (80-) 277:105–109.

35. Braun BR, Head WS, Wang MX, Johnson AD. 2000. Identification and characterization of TUP1-regulated genes in *Candida albicans*. Genetics 156:31–44.

36. Legrand M, Chan CL, Jauert PA, Kirkpatrick DT. 2007. Role of DNA mismatch repair and double-strand break repair in genome stability and antifungal drug resistance in *Candida albicans*. Eukaryot Cell 6:2194–2205.

37. García-Prieto F, Gómez-Raja J, Andaluz E, Calderone R, Larriba G. 2010. Role of the homologous recombination genes RAD51 and RAD59 in the resistance of *Candida albicans* to UV light, radiomimetic and anti-tumor compounds and oxidizing agents. Fungal Genet Biol 47:433–445.

38. Wang G, Tong X, Weng S, Zhou H. 2012. Multiple phosphorylation of Rad9 by CDK is required for DNA damage checkpoint activation. Cell Cycle 11:3792–3800.

39. González-Prieto R, Muñoz-Cabello AM, Cabello-Lobato MJ, Prado F. 2013. Rad51 replication fork recruitment is required for DNA damage tolerance. EMBO J 32:1307–1321.

40. Alcasabas AA, Osborn AJ, Bachant J, Hu F, Werler PJH, Bousset K, Furuya K, Diffley JFX, Carr AM, Elledge SJ. 2001. Mrc1 transduces signals of DNA replication stress to activate Rad53. Nat Cell Biol 3:958–965.

41. Osborn AJ, Elledge SJ. 2003. Mrc1 is a replication fork component whose phosphorylation in response to DNA replication stress activates Rad53. Genes Dev 17:1755–1767.

42. Andaluz E, Ciudad T, Gómez-raja J, Calderone R, Larriba G. 2006. Rad52 depletion in *Candida albicans* triggers both the DNA-damage checkpoint and filamentation accompanied by but independent of expression of hypha-specific genes. Mol mic 59:1452–1472.

43. Lockhart SR, Etienne KA, Vallabhaneni S, Farooqi J, Chowdhary A, Govender NP, Colombo AL, Calvo B, Cuomo CA, Desjardins CA, Berkow EL, Castanheira M, Magobo RE, Jabeen K, Asghar RJ, Meis JF, Jackson B, Chiller T, Litvintseva AP. 2017. Simultaneous emergence of multidrug-resistant *Candida auris* on 3 continents confirmed by whole-genome sequencing and epidemiological analyses. Clin Infect Dis 64:134–140.

44. Chakrabarti A, Sood P, Rudramurthy SM, Chen S, Kaur H, Capoor M, Chhina D, Rao R, Eshwara VK, Xess I, Kindo AJ, Umabala P, Savio J, Patel A, Ray U, Mohan S, Iyer R, Chander J, Arora A, Sardana R, Roy I, Appalaraju B, Sharma A, Shetty A, Khanna N, Marak R, Biswas S, Das S, Harish BN, Joshi S, Mendiratta D. 2014. Incidence, characteristics and outcome of ICU-acquired candidemia in India. Intensive Care Med 41:285–295.

45. Rhodes J, Abdolrasouli A, Farrer RA, Cuomo CA, Aanensen DM, Armstrong-james D, Fisher MC, Schelenz S. 2018. Genomic epidemiology of the UK outbreak of the emerging human fungal pathogen *Candida auris*. Emerg Microbes Infect 7:43:1–12.

46. Zheng X, Wang Y, Wang Y. 2004. Hgc1, a novel hypha-specific G1 cyclin-related protein regulates *Candida albicans* hyphal morphogenesis. EMBO J 23:1845–1856.

47. Braun BR, Kadosh D, Johnson AD. 2001. NRG1, a repressor of filamentous growth in *C. albicans*, is down-regulated during filament induction. EMBO J 20:4753–4761.

48. Kim SH, Iyer KR, Pardeshi L, Muñoz JF, Robbins N, Cuomo CA, Wong KH, Cowen LE. 2019. Genetic analysis of *Candida auris* implicates Hsp90 in morphogenesis and azole tolerance and Cdr1 in azole resistance. MBio 10:e02529–18.

49. Bachewich C, Thomas DY, Whiteway M. 2003. Depletion of a Polo-like kinase in *Candida albicans* activates Cyclase-dependent hyphal-like growth. Mol Biol Evol 14:2163–2180.

50. Chen C, Zeng G, Wang Y. 2018. G1 and S phase arrest in *Candida albicans* induces filamentous growth via distinct mechanisms. Mol Microbiol 110:191–203.

51. Bacal J, Moriel-Carretero M, Pardo B, Barthe A, Sharma S, Chabes A, Lengronne A, Pasero P. 2018. Mrc1 and Rad9 cooperate to regulate initiation and elongation of DNA replication in response to DNA damage. EMBO J 37:1–18.

52. Lopes M, Cotta-Ramusino C, Liberi G, Foiani M. 2003. Branch migrating sister chromatid junctions form at replication origins through Rad51/Rad52-independent mechanisms. Mol Cell 12:1499–1510.

53. Zou L, Elledge SJ. 2003. Sensing DNA damage through ATRIP recognition of RPA-ssDNA complexes. Science (80-) 300:1542–1548.

54. Hashimoto Y, Chaudhuri AR, Lopes M, Costanzo V. 2010. Rad51 protects nascent DNA from Mre11-dependent degradation and promotes continuous DNA synthesis. Nat Struct Mol Biol 17:1305–1311.

55. Petermann E, Orta ML, Issaeva N, Schultz N, Helleday T. 2010. Hydroxyurea-stalled replication forks become progressively inactivated and require two different RAD51-mediated pathways for restart and repair. Mol Cell 37:492–502.

56. Prado F. 2018. Homologous recombination: To fork and beyond. Genes (Basel) 9:603.

57. Zaragoza O, Nielsen K. 2013. Titan cells in *Cryptococcus neoformans*: Cells with a giant impact. Curr Opin Microbiol 16:409–413.

58. Dambuza IM, Drake T, Chapuis A, Zhou X, Correira J, Taylor-Smith L, LeGrave N, Rasmussen T, Fisher MC, Bicanic T, Harrison TS, Jaspars M, May RC, Brown GD, Yuecel R, MacCallum DM, Ballou ER. 2018. The *Cryptococcus neoformans* Titan cell is an inducible and regulated morphotype underlying pathogenesis. PLoS Pathog 14:e1006978.

59. Malavia D, Lehtovirta-Morley LE, Alamir O, Weiß E, Gow NAR, Hube B, Wilson D. 2017. Zinc limitation induces a hyper-adherent Goliath phenotype in *Candida albicans*. Front Microbiol 8:1–14.

60. Loll-krippleber R, Enfert C, Feri A, Diogo D, Perin A, Marcet-houben M, Bougnoux M, Legrand M. 2014. A study of the DNA damage checkpoint in *Candida albicans*: uncoupling of the functions of Rad53 in DNA repair, cell cycle regulation and genotoxic stress-induced polarized growth. Mol Microbiol 91:452–471.

61. Legrand M, Chan CL, Jauert PA, Kirkpatrick DT. 2011. The contribution of the S-phase checkpoint genes MEC1 and SGS1 to genome stability maintenance in *Candida albicans*. Fungal Genet Biol 48:823–830.

62. Katou Y, Kanoh Y, Bando M, Noguchi H, Tanaka H, Ashikari T, Sugimoto K, Shirahige K. 2003. S-phase checkpoint proteins Tof1 and Mrc1 form a stable replication-pausing complex. Nature 424:1078–1083.

63. Masai H, Yang CC, Matsumoto S. 2017. Mrc1/Claspin: a new role for regulation of origin firing. Curr Genet 63:813–818.

64. Miller CT, Gabrielse C, Chen YC, Weinreich M. 2009. Cdc7p-Dbf4p regulates mitotic exit by inhibiting Polo kinase. PLoS Genet 5:e1000498.

65. Asano S, Park JE, Sakchaisri K, Yu LR, Song S, Supavilai P, Veenstra TD, Lee KS. 2005. Concerted mechanism of Swe1/Wee1 regulation by multiple kinases in budding yeast. EMBO J 24:2194–2204.

66. Zhang T, Nirantar S, Lim HH, Sinha I, Surana U. 2009. DNA damage checkpoint maintains Cdh1 in an active state to inhibit anaphase progression. Dev Cell 17:541–551.

67. Simpson-Lavy KJ, Brandeis M. 2010. Clb2 and the APC/CCdh1 regulate Swe1 stability. Cell Cycle 9:3046–3053.

68. David Pruyne and Anthony Bretscher. 2000. Polarization of cell growth in yeast: Establishment and maintenance of polarity states. J Cell Sci 113:365–375.

69. Lew DJ. 2003. The morphogenesis checkpoint: How yeast cells watch their figures. Curr Opin Cell Biol 15:648–653.

70. Ahn SH, Acurio A, Kron SJ. 1999. Regulation of G2/M progression by the STE mitogen-activated protein kinase pathway in budding yeast filamentous growth. Mol Biol Cell 10:3301–3316.

71. Bensen ES, Clemente-Blanco A, Finley KR, Correa-Bordes J, Berman J. 2005. The mitotic cyclins Clb2p and Clb4p affect morphogenesis in Candida albicans. Mol Biol Cell 16:3387–3400.

72. Wu X, Jiang YW. 2005. Possible integration of upstream signals at Cdc42 in filamentous differentiation of *S. cerevisiae*. Yeast 22:1069–1077.

73. Shapiro RS, Cowen LE. 2010. Coupling temperature sensing and development: Hsp90 regulates morphogenetic signaling in *Candida albicans*. Virulence 1:45–48.

74. Khurana N, Laskar S, Bhattacharyya MK, Bhattacharyya S. 2016. Hsp90 induces increased genomic instability toward DNA-damaging agents by tuning down RAD53 transcription. Mol Biol Cell 27:2463–2478.

75. Bravo Ruiz G, Ross ZK, Holmes E, Schelenz S, Gow NAR, Lorenz A. 2019. Rapid and extensive karyotype diversification in haploid clinical *Candida auris* isolates. Curr Genet.

76. Lee KL, Buckley HR, Campbell CC. 1975. An amino acid liquid synthetic medium for the development of mycellal and yeast forms of *Candida albicans*. Sabouraudia 13:148–153.

77. Fortuna M, Sousa MJ, Corte-Real M, Leao C, Salvador A, Sansonetty F. 2000. Cell cycle analysis of yeasts. Curr Protoc Cytom 11.13:1–9.

78. Burke D, Dawson D, Stearns T. 2000. Methods in yeast genetics. Cold Spring Harbor Laboratory.

79. Vyas VK, Barrasa MI, Fink GR. 2015. A *Candida albicans* CRISPR system permits genetic engineering of essential genes and gene families. Sci Adv 1:e1500248.

80. Shevchuk NA, Bryksin A V., Nusinovich YA, Cabello FC, Sutherland M, Ladisch S. 2004. Construction of long DNA molecules using long PCR-based fusion of several fragments simultaneously. Nucleic Acids Res 32:e19.

81. Walther A, Wendland J. 2003. An improved transformation protocol for the human fungal pathogen *Candida albicans*. Curr Genet 42:339–343.

82. Chatterjee S, Alampalli SV, Nageshan RK, Chettiar ST, Joshi S, Tatu US. 2015. Draft genome of a commonly misdiagnosed multidrug resistant pathogen *Candida auris*. BMC Genomics 16:686.

83. Waterhouse AM, Procter JB, Martin DMA, Clamp M, Barton GJ. 2009. Jalview Version 2-a multiple sequence alignment editor and analysis workbench. Bioinformatics 25:1189–1191.

84. Li H, Durbin R. 2010. Fast and accurate long-read alignment with Burrows-Wheeler transform. Bioinformatics 26:589–595.

85. Li H, Handsaker B, Wysoker A, Fennell T, Ruan J, Homer N, Marth G, Abecasis G, Durbin R. 2009. The sequence alignment/Mmp format and SAMtools. Bioinformatics 25:2078–2079.

86. Walker BJ, Abeel T, Shea T, Priest M, Abouelliel A, Sakthikumar S, Cuomo CA, Zeng Q, Wortman J, Young SK, Earl AM. 2014. Pilon: An integrated tool for comprehensive microbial variant detection and genome assembly improvement. PLoS One 9:e112963.

87. Stanke M, Steinkamp R, Waack S, Morgenstern B. 2004. AUGUSTUS: A web server for gene finding in eukaryotes. Nucleic Acids Res 32:309–312.

