## Supplementary Figures & Tables for "Pseudohyphal growth of the emerging pathogen *Candida auris* is triggered by genotoxic stress through the S phase checkpoint"

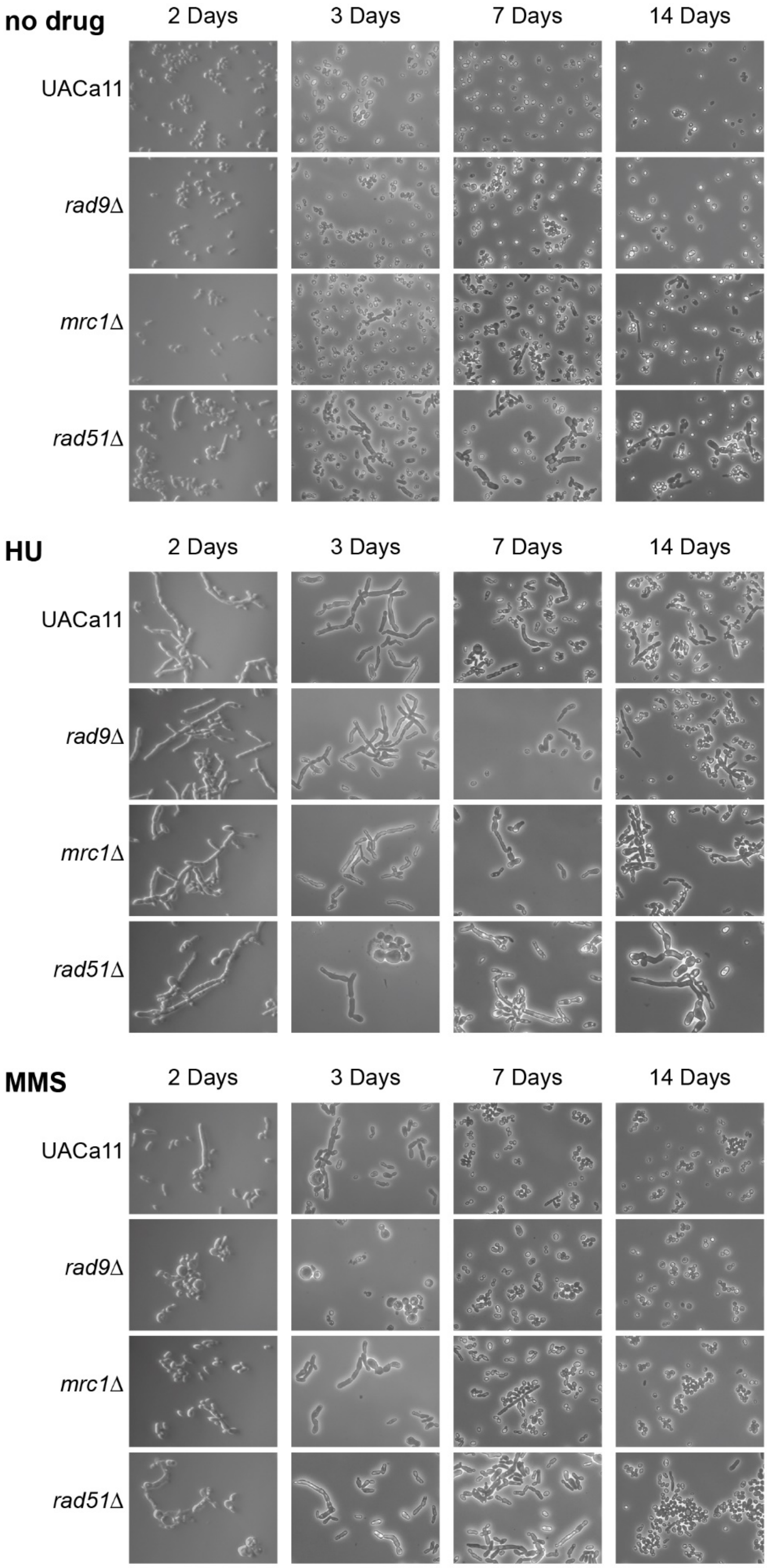


**Figure S1. Filamentation of *Candida auris* wild-type (UACa11) and mutant strains on plates in the presence of genotoxic drugs.** Cells from indicated strains were grown at 30°C on solid YPD media containing no drug, 100 mM Hydroxyurea (HU), or 0.02% methyl methanesulfonate (MMS). Microscopic images were taken at indicated time-points.

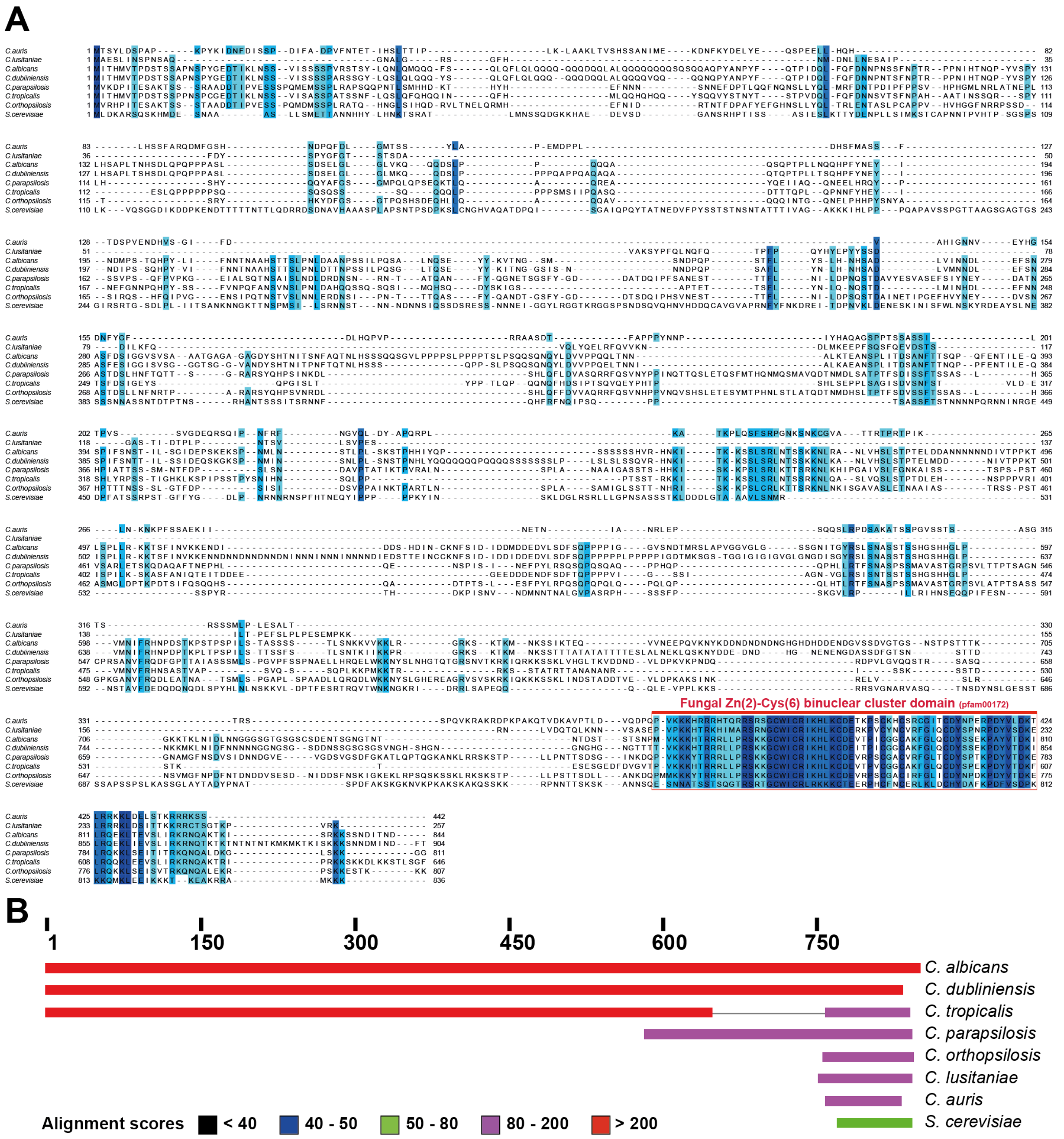


**Figure S2. Alignment of yeast Ume6 protein homologs. (A)** MSAprobs alignment of Ume6 protein sequences from different *Candida* species and *Saccharomyces cerevisiae*. The fungal Zn(2)-Cys(6) domain is highlighted. Shades of blue indicate conservation of >60% of amino acid (characteristics). **(B)** A schematic representation of the alignment scores between yeast Ume6 homologs using the NCBI BLAST-tool (https://blast.ncbi.nlm.nih.gov/Blast.cgi).

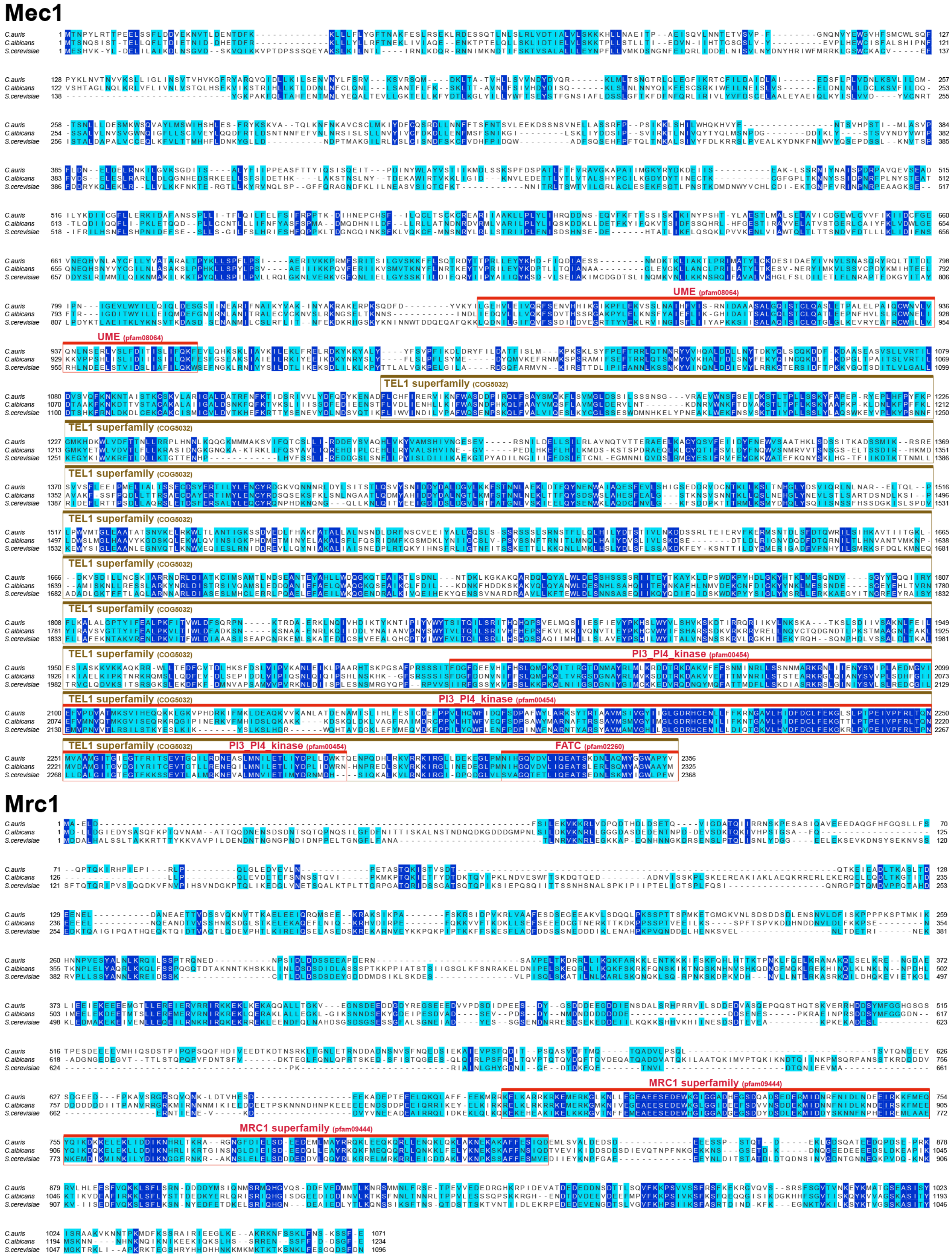


**Figure S3. Alignments of yeast Mec1 and Mrc1 homologs.** MSAprobs alignment of Mec1 and Mrc1 protein sequences from *C. albicans, C. auris* and *S. cerevisiae*. Conserved domains are highlighted. Shades of blue indicate conservation of >60% of amino acid (characteristics).

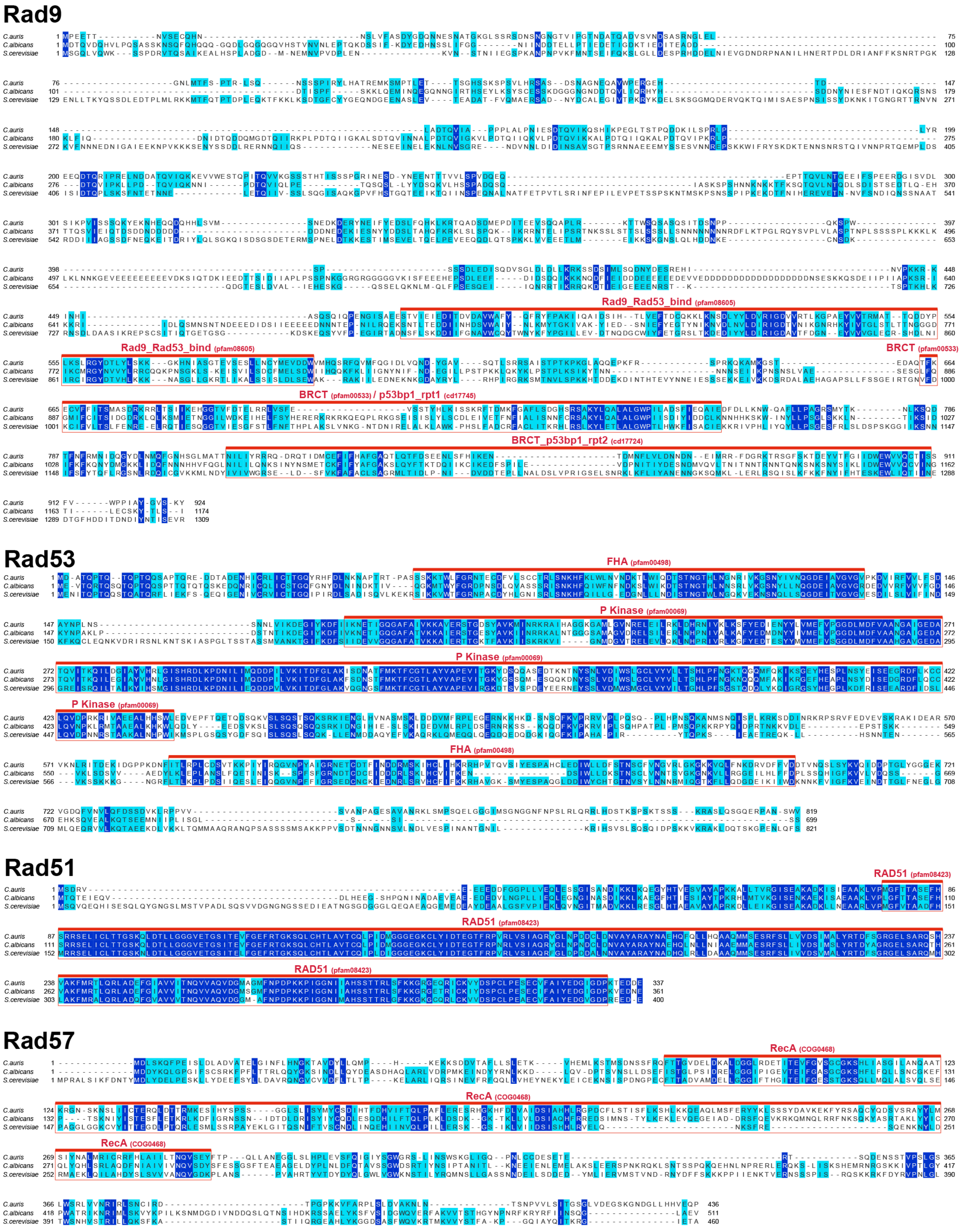


**Figure S4. Alignments of yeast Rad9, Rad53, Rad51 and Rad57 homologs.** MSAprobs alignments of Rad9, Rad53, Rad51 and Rad57 protein sequencess from *C. albicans, C. auris* and *S. cerevisiae*. Conserved domains are highlighted. Shades of blue indicate conservation of >60% of amino acid (characteristics).

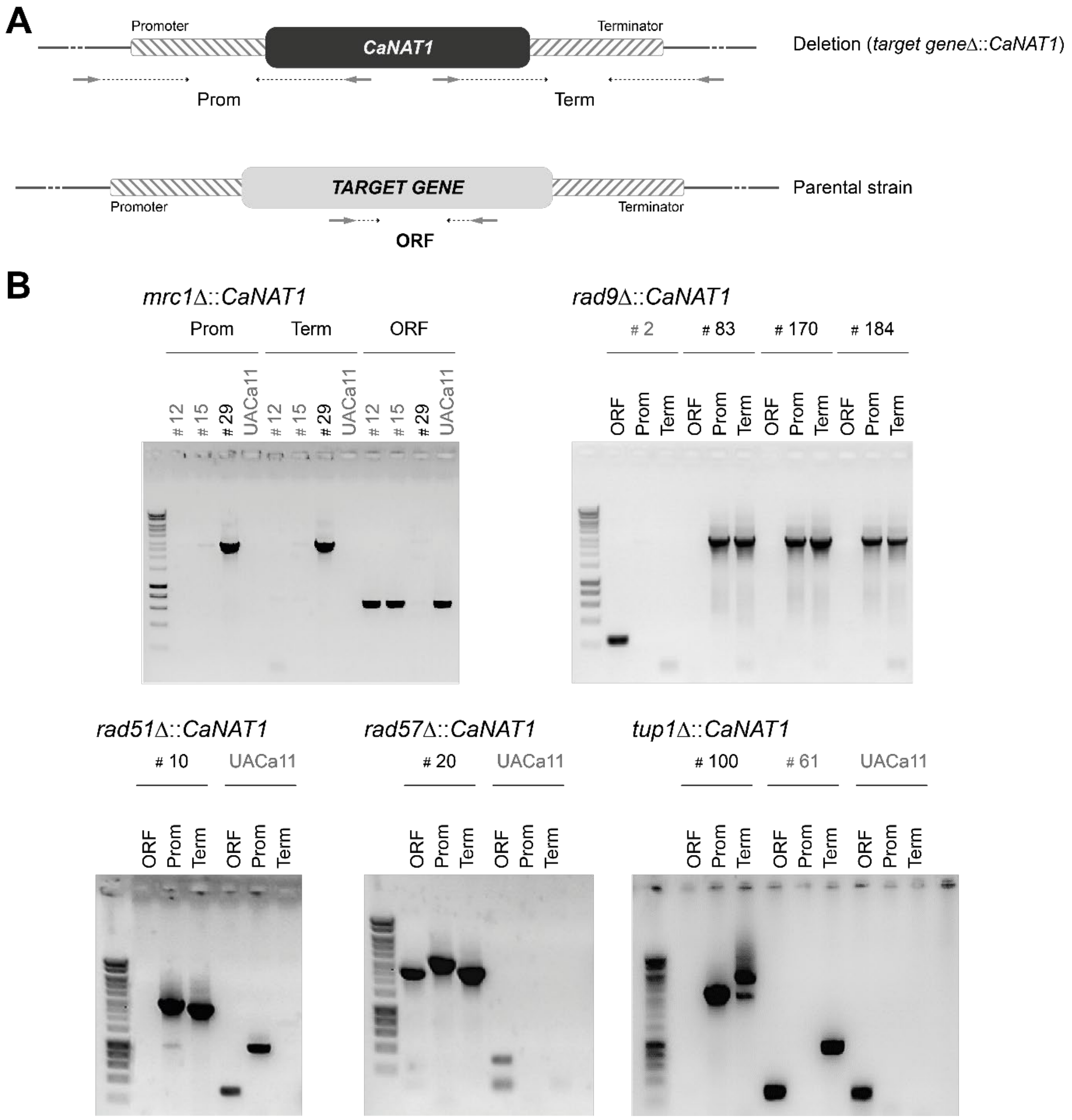


**Figure S5. Verification of *Candida auris* deletion mutants obtained in this study. (A)** Schematic indicating the position of the oligonucleotides used as PCR primers (Table S4) for testing the correct integration of the nourseothricin-resistance marker *CaNAT1* at the target locus in transformants and the absence of the deleted open reading frame (ORF). **(B)** PCR results obtained for indicated deletion strains using the strategy shown in (A). The ORF primers used to verify the *rad57* deletion produced a short unspecific band (<200 bp). Parental wild-type strain UACa11 and transformants with ectopic marker integrations are labelled in light grey. DNA size marker in the left-most lane of every gel is Hyperladder 1kb (Bioline Reagents Ltd, London, UK). Prom = Promoter region; Term = Terminator region.

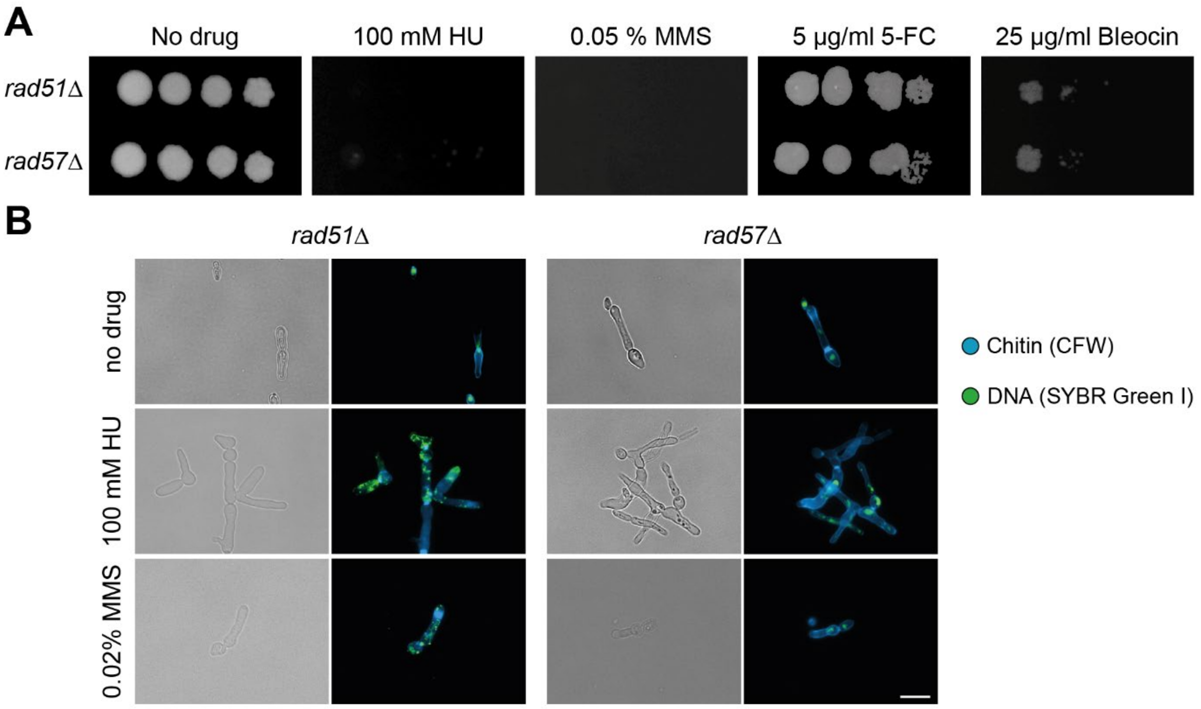


**Figure S6.** ***rad51*** **and** ***rad57*** **deletion strains show similar phenotypes. (A)** Growth analysis of the homologous recombination mutants *rad51*Δ and *rad57*Δ in the presence of genotoxic drugs. Serial dilutions of cells were grown on YPD plates for 3 days at 30°C containing no or the indicated drug; Hydroxyurea (HU), methyl methanesulfonate (MMS), 5-fluorocytosine (5-FC). **(B)** Microscopy images of *rad51*Δ and *rad57*Δ mutant cells grown in YPD broth at 30°C for 18-20 hours containing no or the indicated drug; Hydroxyurea (HU), methyl methanesulfonate (MMS). Chitin was stained with calcofluor white (CFW), and DNA with SYBR Green I. Bright-field images in the left columns and merged fluorescent images in the right columns. Scale bar represents 10 μm.

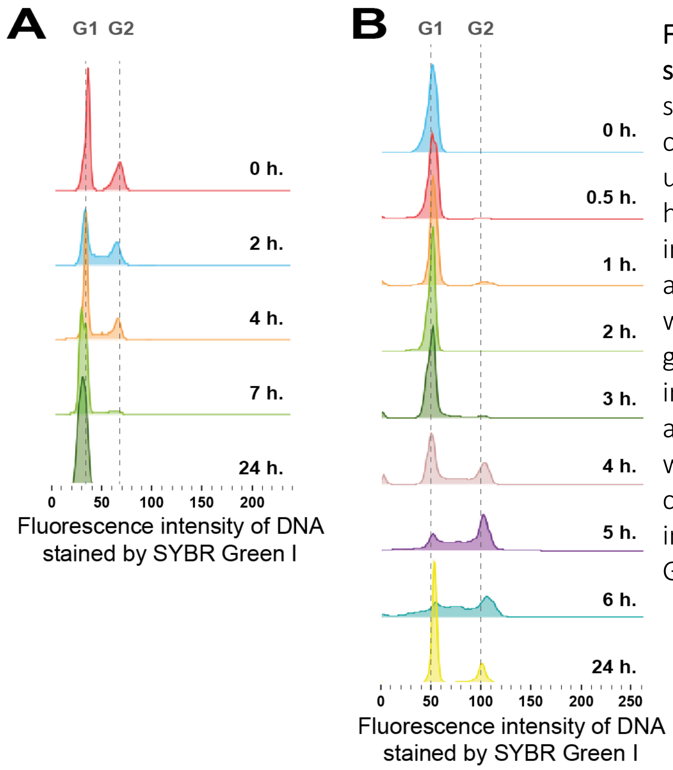


**Figure S7. Cell cycle arrest by nitrogen starvation in *Candida auris*.** Histograms showing cell cycle profiles obtained by flow cytometry of strain UACa11. **(A)** Cells grown under nitrogen starvation conditions for 24 hours at 30°C. Samples were harvested at indicated time points. **(B)** Cells previously arrested by nitrogen starvation for 16 h were inoculated into fresh YPD broth and grown at 30°C. Samples were harvested at indicated time-points. Cells start cycling after a lag phase of 150-180 min. **(A, B)** DNA was stained using SYBR Green I. DNA content is expressed as fluorescence intensity. Approximate position of G1 and G2 peaks are indicated by dotted lines.

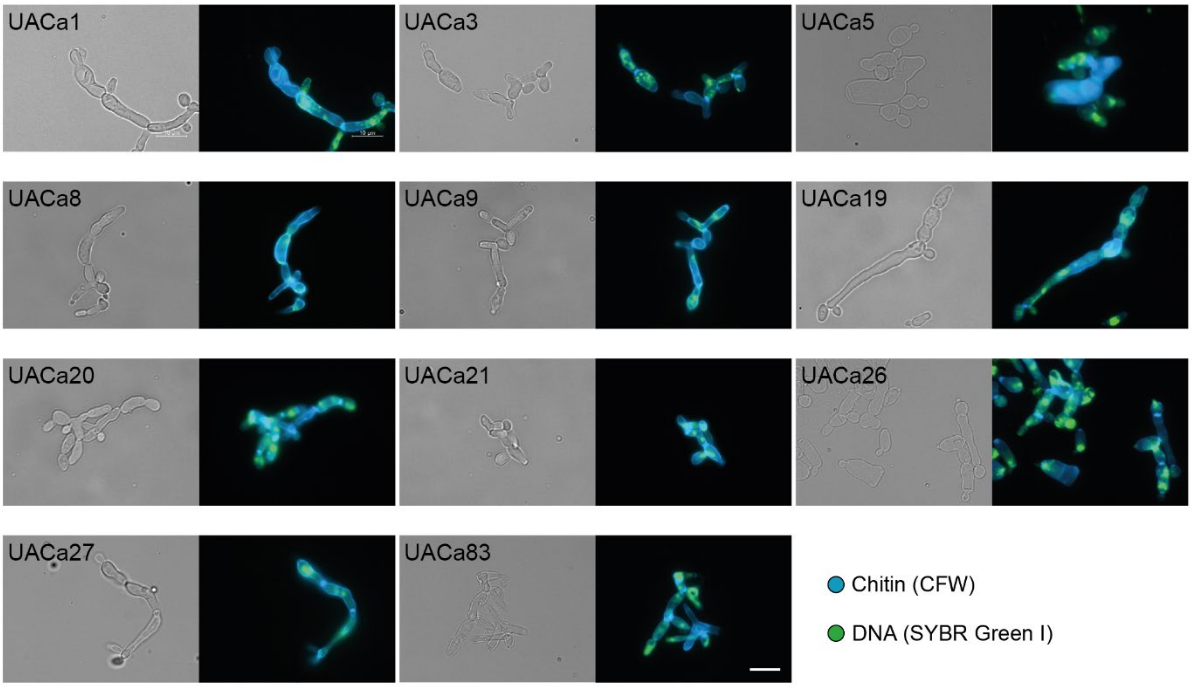


**Figure S8. Different degrees of filamentation in *Candida auris* clinical isolates.** Representative microscopic images of *C. auris* clinical isolates (Table S1) grown for 18-20 hours at 30°C in YPD broth containing 100 mM Hydroxyurea. Chitin was stained with calcofluor white (CFW), and DNA with SYBR Green I. Bright-field images on the left and merged fluorescent images on the right. Scale bar represents 10 μm.

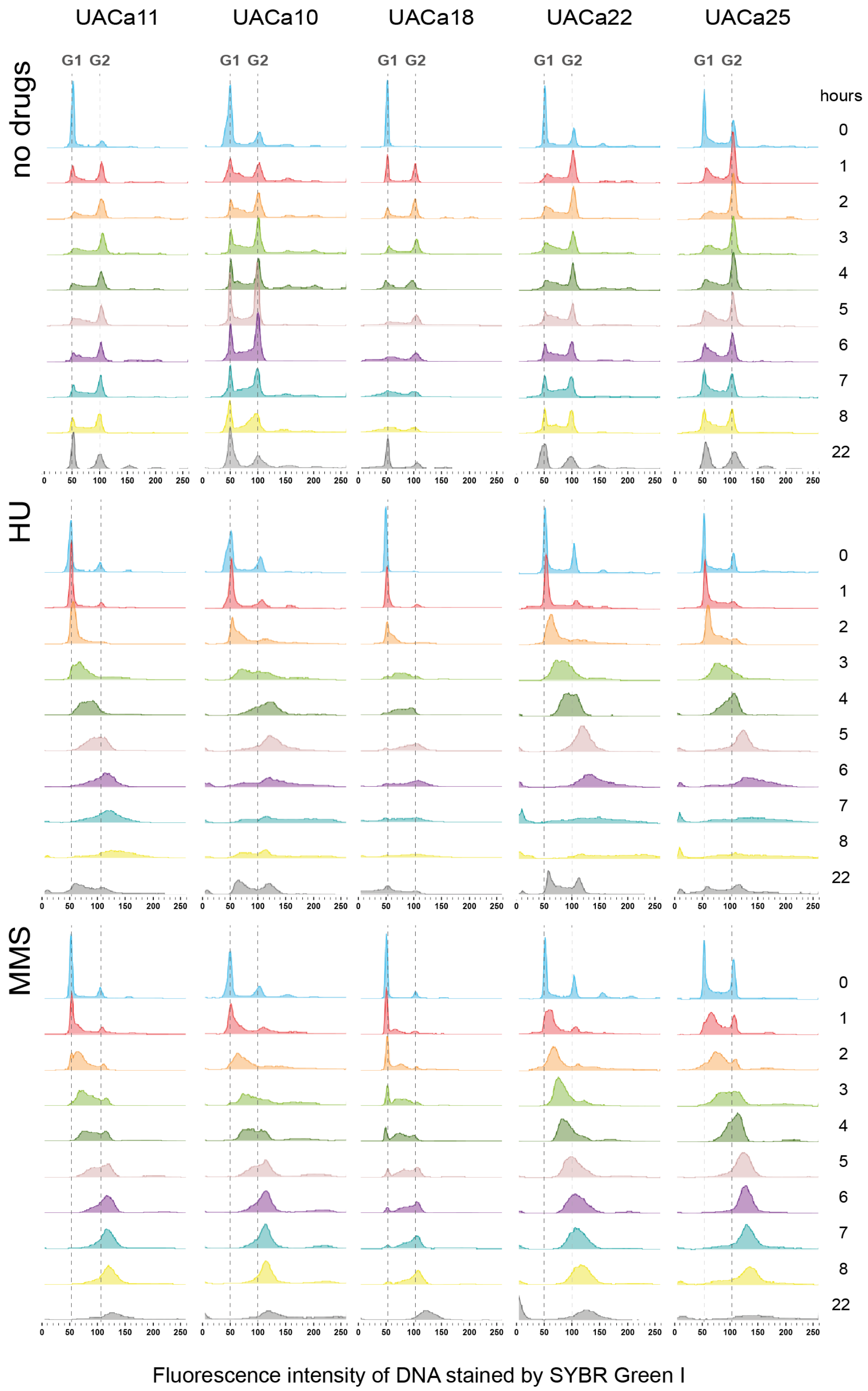


**Figure S9. Cell cycle progression of selected clinical *Candida auris* isolates in the presence of genotoxic drugs.** Histograms showing cell cycle profiles obtained by flow cytometry of indicated clinical isolates. Cells, previously arrested in G1 by nitrogen starvation, were transferred to fresh YPD broth and grown for 165 min at 30°C to restart the cycle before adding 100 mM Hydroxyurea (HU), 0.02% methyl methanesulfonate (MMS), or no drug (time-point 0 hours). Cells were harvested at indicated time-points and DNA stained using SYBR Green I. Amount of DNA is expressed as fluorescence intensity. Approximate positions of G1 and G2 peaks are indicated with dotted lines.

### **Table S1**. Homology among *C. auris*, *C. albicans* and *S. cerevisiae* proteins

| | *C. auris* (Assembly B11221_V1)[I] | | *C. albicans* (SC5314, Assembly 22)[II] | | | | *S. cerevisiae* (S288C)[III] | | | |
|---|---|---|---|---|---|---|---|---|---|---|
| Protein[#] | Access number | Total length | Access number | e-value | Length aligned[†] | Identities (%)[‡] | Access number | e-value | Length aligned[†] | Identities (%)[‡] |
| Ras1 | XP_028890748 | 243 | C2_10210C | 2.00E-77 | 172 | 83.1 | YOR101W | 2.00E-83 | 162 | 77 |
| Ras2 | XP_028892684 | 326 | C3_04480C | 2.00E-19 | 347 | 27.1 | YGR152C | 3.00E-15 | 172 | 28 |
| Nrg1 | XP_028888499 | 255 | C7_04230W | 2.00E-30 | 139 | 53.2 | YDR043C | 4.00E-23 | 57 | 64 |
| Tup1 | XP_028891979 | 601 | C1_00060W | 0 | 406 | 79.1 | YCR084C | 4.00E-180 | 444 | 59 |
| Ume6 | XP_028892977 | 442 | C1_06280C | 2.00E-23 | 74 | 60.8 | YDR207C | 4.00E-11 | 57 | 40 |
| Flo8 | XP_028892793 | 374 | C6_04350C | 2.00E-09 | 93 | 34.4 | YER109C | 6.00E-14 | 118 | 33 |
| Efg1 | XP_028890143 | 495 | CR_07890W | 2.00E-72 | 172 | 76.7 | YMR016C | 3.00E-60 | 108 | 82 |
| Cyr1 | XP_028888959 | 1963 | C7_03070C | 0 | 1765 | 54.4 | YJL005W | 0 | 1421 | 39 |
| Bcy1 | XP_028890128 | 440 | C2_01110C | 6.00E-143 | 450 | 60.9 | YIL033C | 1.00E-127 | 427 | 48 |
| Tpk1/3 | XP_028891209 | 383 | C1_10220C | 0 | 335 | 91.9 | YJL164C (Tpk1) | 0 | 323 | 76 |
| | | | | | | | YKL166C (Tpk3) | 0 | 322 | 77 |
| Tpk2 | XP_028890098 | 448 | C2_07210C | 0 | 357 | 87.7 | YPL203W | 0 | 329 | 85 |
| Eed1 | Not found | | CR_09880W | | | | Not found | | | |
| Cph1 (Ste12) | XP_028890469 | 516 | C1_07370C | 4.00E-97 | 367 | 51.2 | YHR084W | 1.00E-87 | 189 | 66 |
| Cek1 (Fus3) | XP_028892845 | 402 | C4_06480C | 2.00E-168 | 359 | 79.7 | YBL016W | 6.00E-138 | 351 | 58 |
| Cek2 (Kss1) | Not found | | CR_05940W | | | | YGR040W | | | |
| Tec1 | XP_028891285 | 551 | C3_04530C | 8.00E-81 | 380 | 45.3 | YBR083W | 3.00E-16 | 114 | 40 |
| Hgc1 | XP_028888034 | 499 | C1_00780C | 3.00E-86 | 346 | 43.4 | YPL256C | 9.00E-47 | 346 | 30 |
| Flo11 | Not found | | Not found | | | | YIR019C | | | |
| Hwp1 | Not found | | C4_03570W | | | | Not found | | | |
| Hwp2 | Not found | | C4_03510C | | | | Not found | | | |
| Ece1 | Not found | | C4_03470C | | | | Not found | | | |
| Snf1 | XP_028889517 | 604 | C5_01320W | 0 | 578 | 81 | YDR477W | 0 | 598 | 65 |
| Cdc5 | XP_028890727 | 695 | C1_00950C | 0 | 692 | 71.2 | YMR001C | 0 | 637 | 53 |
| Cdc28 | XP_028891635 | 310 | CR_06050W | 8.00E-157 | 292 | 89 | YBR160W | 4.00E-175 | 294 | 76 |
| Clb2 | XP_028891144 | 458 | C2_01410C | 4.00E-150 | 420 | 65.7 | YPR119W | 8.00E-124 | 384 | 50 |
| Gin4 | XP_028888326 | 1270 | C1_11400C | 0 | 1381 | 55.5 | YDR507C | 2.00E-144 | 408 | 58 |
| Hsl7 | XP_028890426 | 480 | Not found | | | | YBR133C | 8.00E-30 | 346 | 29 |
| Hsl1 | XP_028892687 | 1458 | C5_02840C | 0 | 1015 | 47 | YKL101W | 4.00E-125 | 427 | 54 |
| Mih1 | XP_028890158 | 619 | C3_00800W | 4.00E-61 | 385 | 36.9 | YMR036C | 8.00E-37 | 203 | 37 |
| Swe1 | XP_028892733 | 903 | C1_10010C | 1.00E-175 | 1004 | 42.1 | YJL187C | 1.00E-70 | 248 | 48 |
| Rad51 | XP_028892133 | 337 | CR_02200C | 3.00E-176 | 326 | 88 | YER095W | 0 | 330 | 78 |
| Rad53 | XP_028891118 | 819 | C3_03810W | 0 | 690 | 59.3 | YPL153C | 5.00E-158 | 444 | 53 |
| Rad9 | XP_028889586 | 924 | C5_02610C | 4.00E-61 | 500 | 30 | YDR217C | 1.00E-28 | 458 | 24 |
| Mec1 | XP_028890424 | 2356 | C5_04060C | 0 | 2383 | 43.7 | YBR136W | 0 | 1957 | 31 |
| Mrc1 | XP_028891779 | 1071 | C1_11440C | 1.00E-132 | 1233 | 35.5 | YCL061C | 4.00E-32 | 357 | 35 |
| Rad57 | KND99929[$] | 436 | C2_08110W | 3.00E-33 | 259 | 34.4 | YDR004W | 3.00E-16 | 222 | 31 |

[I] *Candida auris* B11221 (UACa20) on NCBI Genome (www.ncbi.nlm.nih.gov/genome/?term=txid498019[orgn])
[II] *Candida* genome database (www.candidagenome.org/)
[III] *Saccharomyces* genome database (www.yeastgenome.org)
[#] *C. albicans*/*S. cerevisiae* names, if different *S. cerevisiae* name in parentheses.
[†] Length of the alignment obtained after BLAST
[‡] Only identities within the alignment obtained
[$] Access number from the *C. auris* Ci6684 draft genome[1].

**Table S2.** Details of *C. auris* strains used in this study

| Strain | Collection No. | Relevant genotype/Clade | Site of isolation | Known drug resistances[a] | Origin/Reference |
|---|---|---|---|---|---|
| 470026 | UACa1 | WT, S. Asia (I) (India) | BSI | FCZ, CSP | A. Chakrabarti[2] |
| 470027 | UACa2 | WT, S. Asia (I) (India) | BSI | FCZ, VCZ, CSP | A. Chakrabarti[2] |
| 470028 | UACa3 | WT, S. Asia (I) (India) | BSI | FCZ, CSP | A. Chakrabarti[2] |
| 470029 | UACa4 | WT, S. Asia (I) (India) | BSI | FCZ, VCZ, CSP | A. Chakrabarti[2] |
| 470030 | UACa5 | WT, S. Asia (I) (India) | BSI | FCZ, VCZ, CSP | A. Chakrabarti[2] |
| NCPF8980#9 | UACa6 | WT, S. Africa (III) | BSI | FCZ, CSP | E. Johnson |
| NCPF8984#15 | UACa7 | WT, E. Asia (II) (Japan) | unknown | FCZ | E. Johnson |
| NCPF8985#20 | UACa8 | WT, S. Asia (I) (India) | wound | FCZ, ISA, PSZ, VCZ, 5-FC, AFG | E. Johnson |
| NCPF13001#16 | UACa9 | WT, S. Asia (I) (India) | unknown | FCZ | E. Johnson |
| NCPF13005#95 | UACa10 | WT, S. Africa (III) | urine | FCZ, VCZ, AFG, AMB, CSP | E. Johnson |
| VPCI479/P/13 | UACa11 | WT, S. Asia (I) (India) | BSI | FCZ | A. Chowdhary[3] |
| B11220 | UACa18 | WT, E. Asia (II) (Japan) | auditory canal | FCZ, VCZ | S. Lockhart[4] |
| B11109 | UACa19 | WT, S. Asia (I) (Pakistan) | burn wound | CSP | S. Lockhart[4] |
| B11221 | UACa20 | WT, S. Africa (III) | BSI | FCZ, CSP | S. Lockhart[4] |
| B11222 | UACa21 | WT, S. Africa (III) | BSI | FCZ, CSP | S. Lockhart[4] |
| B11244 | UACa22 | WT, S. America (IV) (Venezuela) | BSI | FCZ, VCZ, CSP | S. Lockhart[4] |
| B11245 | UACa23 | WT, S. America (IV) (Venezuela) | BSI | FCZ, VCZ, CSP | S. Lockhart[4] |
| B8441 | UACa24 | WT, S. Asia (I) (Pakistan) | BSI | CSP | S. Lockhart[4] |
| B11098 | UACa25 | WT, S. Asia (I) (Pakistan) | BSI | FCZ | S. Lockhart[4] |
| B11203 | UACa26 | WT, S. Asia (I) (India) | BAL | FCZ, 5-FC | S. Lockhart[4] |
| B11205 | UACa27 | WT, S. Asia (I) (India) | chest wound | FCZ, VCZ, 5-FC | S. Lockhart[4] |
| CBS10913T | UACa83 | WT, E. Asia (I) (Japan) | auditory canal | none | CBS-KNAW collection[5] |
| | UACa93 | *rad51*Δ::*CaNAT1* (UACa11 BCKG) | lab strain | not tested | this study |
| | UACa94 | *rad57*Δ::*CaNAT1* (UACa11 BCKG) | lab strain | not tested | this study |
| | UACa95 | *tup1*Δ::*CaNAT1* (UACa11 BCKG) | lab strain | not tested | this study |
| | UACa109 | *rad9*Δ::*CaNAT1* (UACa11 BCKG) | lab strain | not tested | this study |
| | UACa112 | *mrc1*Δ::*CaNAT1* (UACa11 BCKG) | lab strain | not tested | this study |

[a]Using EUCAST clinical breakpoints for *Candida albicans* (http://www.eucast.org/fileadmin/src/media/PDFs/EUCAST_files/AFST/Clinical_breakpoints/Antifungal_breakpoints_v_9.0_180212.pdf)

Abbreviations: BAL = broncho-alveolar lavage, BSI = bloodstream infection, FCZ = fluconazole, ISA = Isavuconazole, PSZ = Posaconazole, VCZ = Voriconazole, CSP = Caspofungin, AFG = Anidulafungin, 5-FC = flucytosine, AMB = Amphotericin B, WT = Wild Type, BCKG = background.

**Table S3.** Non-synonymous SNPs between UACa1 and UACa4

| Protein[1] | Description | SNP[2] |
|---|---|---|
| Cdc37p | Chaperone for Crk1 (involved in hyphal development of Cph1- and Efg1-independent) | Ser-46-Asn |
| XP_028889655[3] | Hypothetical protein, fungal specific Zn(II)2Cys6 transcription factor, possibly related to *C. albicans* Zcf32 | Arg-458-STOP |
| Kin2p | Hyphal growth, cell growth switching | Asp-582-Glu |
| Tpk2p | cAMP-dependent protein kinase catalytic subunit; isoform of Tpk1; regulation of filamentation, phenotypic switching and mating | Asp-32-Glu |
| Hap1p | Predicted Zn(II)2Cys6 transcription factor, filamentous growth in response to biotic stimulus | Pro-967-Ala |
| Cdc3p | Septin; hyphal growth; macrophage/pseudohyphal-repressed | His-214-Pro |
| Mds3p | TOR signalling pathway component; required for growth and hyphal formation at alkaline pH; pseudohyphal growth | Leu-437-Ser |
| Dot1p | White-opaque switching; DNA damage checkpoint | Ser-641-Thr |
| Isw2p | ATPase involved in chromatin remodeling; Hap43-induced gene; Filamentous growth | Lys-462-Met |
| Dyn1p | Motor protein that moves to microtubule minus end; yeast cell separation, spindle positioning, nuclear migration, hyphal growth | Gly-1995-Arg |
| Sch9p | Protein kinase; involved in growth control, cell size, filamentous growth under some conditions, and virulence | Ile-544-Leu |
| Mss11p | Transcription factor; activator that binds to Flo8; required for hyphal growth | Gly-134-Arg |
| Arc40p | Involved in actin filament organization; affects filamentous growth | His-91-Asn |
| Gpr1p | Plasma membrane G-protein-coupled receptor of the cAMP-PKA pathway; required for hyphal growth | Gly-167-Ser |
| Wor2p | Regulator of white-opaque switching; filamentous growth | Phe-91-Leu |
| Tel1p | Telomeric DNA binding activity-DNA repair/damage checkpoint | Ala-2001-Thr |
| Tif4631p | Translation initiation factor eIF4G; overexpression causes hyperfilamentation; hyphal- and macrophage-induced | Asp-763-Gly |
| Gin4p | Autophosphorylated kinase; role in pseudohyphal-hyphal switch and cytokinesis | Ser-507-Asn |
| Ecm29p | Scaffold protein; association of the proteasome core particle with the regulatory particle; mutation affects filamentous growth | Ser-489-Tyr |
| Hal9p | Zn(II)2Cys6 transcription factor involved in salt tolerance, Filamentous growth | Ala-640-Val |
| Hap1p | Predicted Zn(II)2Cys6 transcription factor, filamentous growth in response to biotic stimulus | Asn-524-His |
| Tip41p | TOR signalling pathway; regulates Rad53p during DNA damage, Regulation filamentous growth | Phe-331-Leu |
| YPL108W-like protein | YPL108W is a non-essential gene induced in a GDH1 deleted strain; induced in response to MMS | Gln-92-STOP |
| Vps36p | ESCRT II protein sorting complex subunit; regulation of pH response, filamentation in response to pH | Ala-441-Thr |

[1]*S. cerevisiae* protein names, unless otherwise indicated.
[2]Amino acid produced in UACa1 in front of and in UACa4 after indicated position, stop codon indicated as STOP.
[3]Systematic ID of *Candida auris* B11221 (UACa20) on NCBI Genome (www.ncbi.nlm.nih.gov/genome/?term=txid498019[orgn])

**Table S4.** Primers used in this study

| Name | sequence 5'-3'* | Orientation | Experimental use |
|---|---|---|---|
| ***CaNAT1*** | | | |
| oUA315 | tcgtacgctgcaggtcgacgGTCGACACTGGATGGCGG | forward | for pALo218 construction (pFA6a overhang) |
| oUA316 | cgcgccttaattaacccgggATCAAGCTTGCCTCGTCC | reverse | for pALo218 construction (pFA6a overhang) |
| oUA345 | GCCGCGCCCCTGTAGAGAAA | reverse | ORF for mutants check |
| oUA346 | CTCTGGCGAATTCAGTAGCCAAACC | forward | ORF for mutants check |
| oUA353 | ATCAAGCTTGCCTCGTCCCC | forward | for gene deletion construction |
| oUA354 | GTCGACACTGGATGGCGGC | reverse | for gene deletion construction |
| ***RAD51*** | | | |
| oUA570 | tgctgtcgattcgatactaacgccgccatccagtgtcgacATAACGCGCCACCCAAAGAG | reverse | promoter region for gene deletion construction (CaNAT1 overhang) |
| oUA571 | GCTGCTTGACTCGGTTCCAC | forward | promoter region for gene deletion construction / mutants check |
| oUA572 | CGTCTTTGAGTCGATGTGTTGGATG | forward | promoter region nested primer for whole construction amplification |
| oUA573 | cgctggccgggtgacccggcggggacgaggcaagcttgatGGGCTTTATCCAAAGTCAGGGAA | forward | terminator region for gene deletion construction (CaNAT1 overhang) |
| oUA574 | CGGCAAATTGATCAATGGCAGAG | reverse | terminator region for gene deletion construction / mutants check |
| oUA575 | CCATGCAGCATGAAATGGTTATGC | reverse | terminator region nested primer for whole construction amplification |
| oUA576 | AAACTCGTGCCTATGGGCTTCA | forward | ORF for mutants check |
| oUA577 | AGTCTCACGGGTCTGAATGTGC | reverse | ORF for mutants check |
| ***RAD57*** | | | |
| oUA578 | tgctgtcgattcgatactaacgccgccatccagtgtcgacTTCGGTTTCACTCGATGCGAAAC | reverse | promoter region for gene deletion construction (CaNAT1 overhang) |

| | | | |
|---|---|---|---|
| oUA579 | CTCGGTTGACGACAGAGACTATGG | forward | promoter region for gene deletion construction / mutants check |
| oUA580 | CTCAGAGAGTCGCTATCAAAGTCGA | forward | promoter region nested primer for whole construction amplification |
| oUA581 | cgctggccgggtgacccggcggggacgaggcaagcttgatTTCATGCAGCTCTCGCGTCT | forward | terminator region for gene deletion construction (CaNAT1 overhang) |
| oUA582 | ACTGATCGCCATCCCACCTAAATC | reverse | terminator region for gene deletion construction / mutants check |
| oUA583 | CTGTGCGGGGGCTACAATTAGA | reverse | terminator region nested primer for whole construction amplification |
| oUA584 | TCGGTGTAAGTGGATGTGGCAA | forward | ORF for mutants check |
| oUA585 | AAAGTGCTTTCCATGGCGGC | reverse | ORF for mutants check |
| ***RAD9*** | | | |
| oUA624 | tgctgtcgattcgatactaacgccgccatccagtgtcgacTCGATACTGGAATAGTGGTGGTTGG | reverse | promoter region for gene deletion construction (CaNAT1 overhang) |
| oUA625 | GAAGCTGTGGCAAGCAAGCTT | forward | promoter region for gene deletion construction / mutants check |
| oUA626 | GGCTCGTTATCATTTCTGGATGCC | forward | promoter region nested primer for whole construction amplification |
| oUA627 | cgctggccgggtgacccggcggggacgaggcaagcttgatGCAAAGAGCATTCTATGCGAAGAGC | forward | terminator region for gene deletion construction (CaNAT1 overhang) |
| oUA628 | GCTCATCACATTTCTTTTCGAGGGG | reverse | terminator region for gene deletion construction / mutants check |
| oUA629 | CAAGTCAGTTGATCTCTCTCCTGCA | reverse | terminator region nested primer for whole construction amplification |
| oUA676 | CAACGAATCGAATGCAACCGGG | forward | ORF for mutants check |
| oUA677 | GACGAGGACGGACTCCATGG | reverse | ORF for mutants check |
| ***MRC1*** | | | |
| oUA703 | CTCCACCATGTCAGCATACCACA | forward | promoter region for gene deletion construction / mutants check |
| oUA704 | GACGAAATCCAACCTGGGGCT | forward | promoter region nested primer for whole construction amplification |
| oUA705 | cgctggccgggtgacccggcggggacgaggcaagcttgatTGAACAGCAGCTGGCTCCTA | forward | terminator region for gene deletion construction (CaNAT1 overhang) |

| | | | |
|---|---|---|---|
| oUA706 | CCAAACAGGTGAATGTGCAGCT | reverse | terminator region for gene deletion construction / mutants check |
| oUA707 | AGCGAGCGAGTGTCTCAACTTT | reverse | terminator region nested primer for whole construction amplification |
| oUA708 | CTACTGTGGACTCGTCCGTGC | forward | ORF for mutants check |
| oUA709 | TTCAGCATCTCCGTTTTCACGC | reverse | ORF for mutants check |
| oUA778 | tgctgtcgattcgatactaacgccgccatccagtgtcgacGAAGTGTTAGCAATTGCTGGAGAG | reverse | promoter region for gene deletion construction (CaNAT1 overhang) |
| ***TUP1*** | | | |
| oUA588 | tgctgtcgattcgatactaacgccgccatccagtgtcgacGTGGATGGGGCACAGAATTAGAGA | reverse | promoter region for gene deletion construction (CaNAT1 overhang) |
| oUA589 | CCGACTCACCAGAATATAGCGGTC | forward | promoter region for gene deletion construction / mutants check |
| oUA590 | GCAGGTAAATGCGGGCAAAGG | forward | promoter region nested primer for whole construction amplification |
| oUA591 | cgctggccgggtgacccggcggggacgaggcaagcttgatCAGTTTTCATGAGCATGGGGTCG | forward | terminator region for gene deletion construction (CaNAT1 overhang) |
| oUA592 | GTATGAGCGTTGCTGCATACCAG | reverse | terminator region for gene deletion construction / mutants check |
| oUA593 | GTGCCACAGTGCAAACTGTCATC | reverse | terminator region nested primer for whole construction amplification |
| oUA594 | GGACCCATCGAAAGCTCTGGA | forward | ORF for mutants check |
| oUA595 | CCAAAGATTGGACCAAGTCCACG | reverse | ORF for mutants check |

*Overhangs in lowercase
ORF = Open reading frame

### Supplementary References

1. Chatterjee S, Alampalli SV, Nageshan RK *et al.* Draft genome of a commonly misdiagnosed multidrug resistant pathogen *Candida auris*. *BMC Genomics* 2015;**16**:686.

2. Chakrabarti A, Sood P, Rudramurthy SM *et al.* Incidence, characteristics and outcome of ICU-acquired candidemia in India. *Intensive Care Med* 2014;**41**:285–95.

3. Sharma C, Kumar N, Meis JF *et al.* Draft genome sequence of a fluconazole-resistant *Candida auris* strain from a candidemia patient in India. *Genome Announc* 2015;**3**:e00722-15.

4. Lockhart SR, Etienne KA, Vallabhaneni S *et al.* Simultaneous emergence of multidrug-resistant *Candida auris* on 3 continents confirmed by whole-genome sequencing and epidemiological analyses. *Clin Infect Dis* 2017;**64**:134–40.

5. Satoh K, Makimura K, Hasumi Y *et al. Candida auris* sp. nov., a novel ascomycetous yeast isolated from the external ear canal of an inpatient in a Japanese hospital. *Microbiol Immunol* 2009;**53**:41–4.
